# FAM114A1 Influences Cardiac Fibrosis by Regulating Angiotensin II Signaling in Cardiac Fibroblasts

**DOI:** 10.1101/2021.02.20.432115

**Authors:** Kadiam C Venkata Subbaiah, Jiangbin Wu, Wai Hong Wilson Tang, Peng Yao

## Abstract

Cardiac fibrosis, a primary contributor to heart failure (HF) and sudden death, is considered as an important target for HF therapy. However, the signaling pathways that govern cardiac fibroblast (CF) function during cardiac fibrosis have not been fully elucidated. Here, we found that a functionally unannotated human myocardial infarction (MI) associated gene, family with sequence similarity 114 member A1 (FAM114A1), is induced in failing human and mouse hearts compared to non-failing hearts. Homozygous knockout of *Fam114a1* (*Fam114a1*^−/−^) in the mouse genome reduces cardiac hypertrophy and fibrosis while significantly restores cardiac function in angiotensin (Ang) II- and MI-induced HF mouse models. *Fam114a1* deletion antagonizes Ang II induced inflammation and oxidative stress. Using isolated mouse primary CFs in wild type and *Fam114a1*^−/−^ mice, we found that FAM114A1 is a critical autonomous factor for CF proliferation, activation, and migration. We discovered that FAM114A1 interacts with angiotensin receptor associated protein (AGTRAP) and regulates the expression of angiotensin type 1 receptor (AT1R) and downstream Ang II signaling transduction, and subsequently influences pro-fibrotic response. Using RNA-Seq in mouse primary CFs, we identified differentially expressed genes including extracellular matrix proteins such as Adamts15. RNAi-mediated inactivation of Adamts15 attenuates CF activation and collagen deposition. Our results indicate that FAM114A1 regulates Ang II signaling and downstream pro-fibrotic and pro-inflammatory gene expression, thereby activating cardiac fibroblasts and augmenting pathological cardiac remodeling. These findings provide novel insights into regulation of cardiac fibrosis and identify FAM114A1 as a new therapeutic target for treatment of cardiac disease.

**Significance:** Cardiac fibrosis is a hallmark of heart failure and angiotensin II signaling promotes pro-fibrotic response in the heart. This study is a pioneering investigation of the role of a functionally unknown protein FAM114A1. We show that FAM114A1 expression is induced in human and mouse failing hearts. Genetic ablation of FAM114A1 can effectively reduce cardiac fibrosis and pathological remodeling. Isolated cardiac fibroblasts from *Fam114a1* knockout mice show reduced response to Ang II stimulation and compromised myofibroblast activation. Mechanistically, FAM114A1 binds to AGTRAP and influences AT1R protein expression, thereby enhancing angiotensin II signaling and pro-fibrotic response. Thus, FAM114A1 is a novel factor that modulates cardiac fibrosis and pharmacological inhibition of FAM114A1 may be a therapeutic strategy for the treatment of heart disease.

## Introduction

Heart failure (HF) is a leading cause of morbidity and mortality across the globe (1). The development of HF in the context of chronic stresses such as hypertension and myocardial infarction (MI) is characterized by complex changes in the structure and function of the heart at the cellular and molecular levels. This cardiac pathological remodeling process ultimately leads to ventricular hypertrophy, chamber dilatation, contractile dysfunction, and HF. Furthermore, massive clinical and experimental data have demonstrated that chronic activation of cardiac fibroblasts (CFs) and accumulation of cardiac fibrosis adversely affect cardiac compliance, cause diastolic dysfunction (2, 3), and leads to arrhythmia and sudden death (4, 5). Even though cardiac fibrosis is clinically important in HF, effective anti-fibrosis therapeutics are not available till date (6, 7). Therefore, understanding the underpinning mechanisms of cardiac fibrosis and identifying novel modalities to modulate this process are of high scientific impact and therapeutic potential.

In response to cardiac injuries during pathological remodeling, quiescent CFs undergo a remarkable cell state transition into myofibroblasts (MFs) with increased collagen production, proliferation, and contractility (8), thereby driving cardiac fibrosis (9). At the molecular level, angiotensin II (Ang II), a key neurohormonal ligand constituent of the renin-angiotensin system (RAS), plays a vital role in cardiac fibrosis via promoting CF proliferation and increasing the production of pro-inflammatory cytokines and extracellular matrix (ECM) proteins through the activation of angiotensin II type-1 receptor (AT1R) (10, 11). The function of circulating RAS has been extensively studied. What remain elusive is the regulatory mechanism of tissue- and cell type-specific RASs that are involved in long-term effects of chronic RAS activation associated with local organ damage in the cardiac pathophysiology (12). Ang II activates pro-fibrogenic cascades such as TGF*β*/SMAD (transforming growth factor beta), ERK1/2 (extracellular signal-regulated kinases 1 and 2), AKT (protein kinase B), and p38 MAPK (mitogen-activated protein kinase) signal transduction during the pathogenesis of cardiac fibrosis (13-16). The pathophysiological effects of Ang II can rewire the transcriptome of CFs to promote the activation of fibroblasts with enhanced ability to proliferate, migrate, secrete pro-inflammatory mediators, and produce ECM proteins (10, 11, 13, 14). Indeed, the first-line treatment for HF relies on angiotensin-converting enzyme (ACE) inhibitors and Ang II receptor blockers, which show a promise in blunting cardiac fibrosis (2, 6, 17). This treatment, however, is limited due to its adverse side effects. Thus, we still need to identify cell type-specific Ang II signaling regulators to advance the therapeutic interventions. Since CFs are critical players in the cardiac pathogenesis and HF, targeting the Ang II signaling cascade in CFs is of utmost importance for the development of effective therapies against HF (18).

In Ang II signaling transduction, AT1R serves as a control point for regulating the Ang II effects and AT1R over-activation leads to cardiac pathogenesis (19, 20). Angiotensin II type 1 receptor-associated protein (AGTRAP) selectively inhibits AT1R over-activation and suppresses the Ang II-induced cardiac remodeling in mouse HF models (21, 22). CM-specific AGTRAP transgenic mice exhibit significantly diminished Ang II-induced cardiac remodeling due to increased AGTRAP:AT1R ratio, AT1R internalization, and diminished AT1R downstream effects (23). Interestingly, AGTRAP expression was significantly reduced in mouse failing hearts (23). However, the molecular mechanism is poorly understood (11). A yeast two-hybrid screen identified AGTRAP as an interacting protein of family with sequence similarity 114 member A1 (FAM114A1) (24). However, the function and mechanism of FAM114A1 in cardiac pathophysiology and general biology is unknown to date. Therefore, we investigated the role of FAM114A1 in heart disease and its potential as a therapeutic target for HF treatment.

This work has revealed the role of FAM114A1 in CFs and during cardiac fibrosis. FAM114A1 expression is robustly elevated in human and mouse failing hearts, highly enriched in CFs, and correlated with the expression of pro-fibrotic collagens. *Fam114a1*^−/−^ mouse hearts display greater resistance to Ang II- and MI-induced cardiac inflammation, fibrosis and dysfunction compared to hearts from wild type mice. Mechanistically, FAM114A1 binds to AGTRAP and increases AT1R expression, thereby amplifying Ang II signaling. Transcriptomic profiling of *Fam114a1* null CFs revealed multiple downstream effectors involved in ECM remodeling, such as Adamts15 that is required for cardiac fibroblast activation and migration.

## Results

### FAM114A1 expression is induced in cardiac fibroblasts of human and mouse failing hearts

Three human genome-wide association studies have discovered that a single nucleotide polymorphism (SNP) rs1873197 in *FAM114A1* genomic locus is associated with MI and coronary artery disease (CAD) (25-27). These genetic associations of *FAM114A1* gene with MI and CAD imply its potential impact in cardiac pathogenesis. From data mining of high throughput screen for upregulated miRNAs in cardiac pathogenesis, we identified miR-574 as a miRNA that is consistently induced in the human and mouse heart in response to pathological cardiac remodeling (28-31). The host gene of miR-574, *FAM114A1*, has been shown to be co-expressed with the embedded intronic miRNA in the brain (32). Therefore, we attempted to examine whether FAM114A1 expression is also induced by cardiac stresses. Using human HF patient samples, we found that FAM114A1 was induced by ∼50% at both mRNA and protein levels in human failing hearts, compared to non-failure donor hearts (Figure 1A, B). To explore the gene expression of FAM114A1 in mouse HF, we infused the wild type (WT) mice with Ang II (1.4 mg/Kg/day) for 4 weeks to mimic human hypertensive cardiomyopathy. Immunoblotting and RT-qPCR showed that Ang II induced the expression of FAM114A1 by ∼4-fold at both mRNA and protein levels in murine hearts (Figure 1C, D).

**Figure 1.**
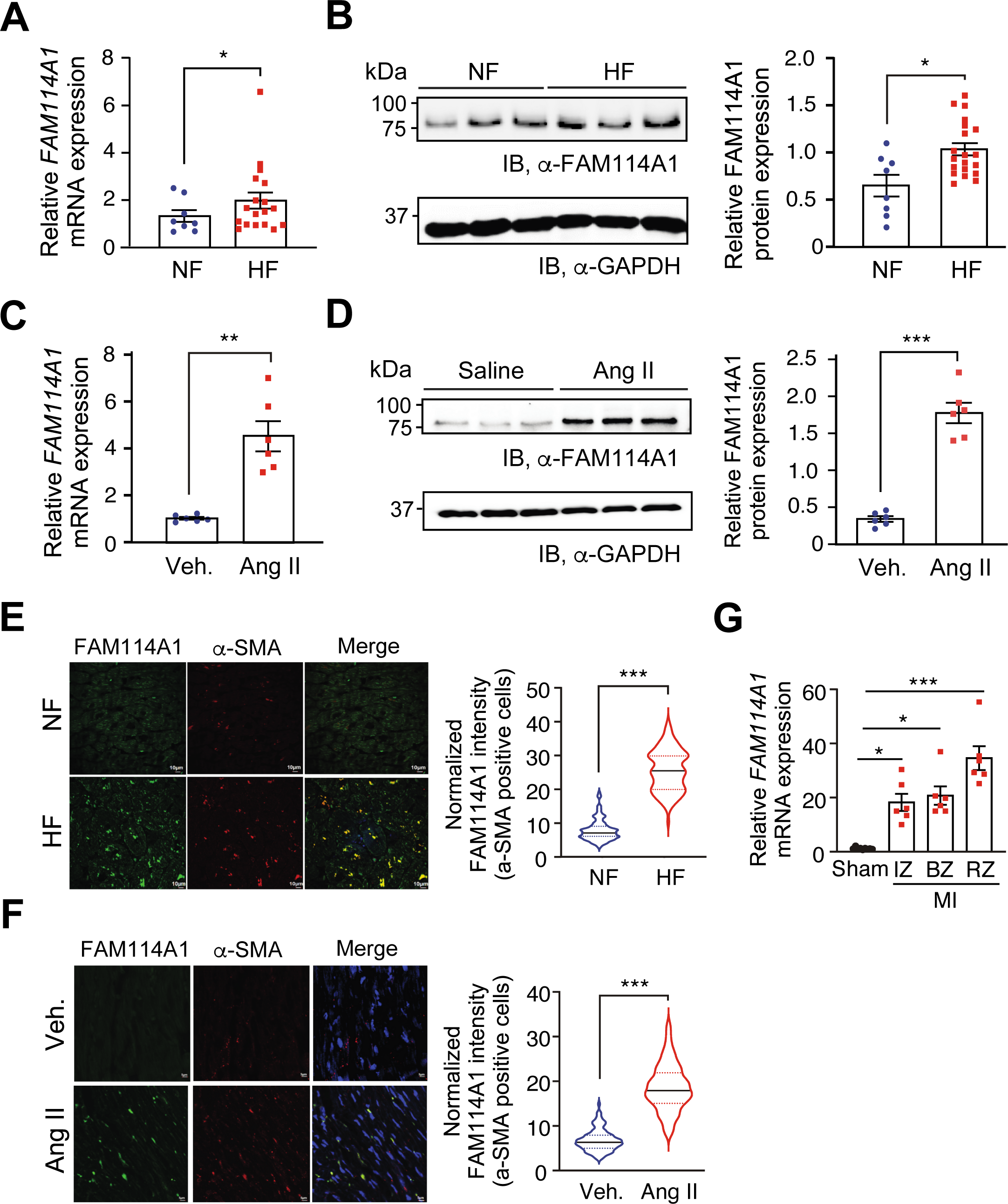
Increased FAM114A1 expression in activated myofibroblasts in human and mouse failing heart. (A) *FAM114A1* mRNA expression is increased in failing human hearts (n=18) compared to non-failure donor hearts (n=8). 18S rRNA was used as a normalizer. (B) FAM114A1 protein expression is increased in failing human hearts (n=20) compared to non-failure hearts (n=8). GAPDH protein was used as a normalizer. (C-D) FAM114A1 mRNA and protein expression is significantly induced in the hearts from mice with Ang II infusion (4 weeks) compared to hearts from vehicle treated mice (n=6 for both groups). 18S rRNA and *Gapdh* mRNA were used as normalizers for mRNA and protein quantification, respectively. (E-F) Immunofluorescence analysis and quantification of FAM114A1 protein expression in human failing hearts and Ang II treated mouse heart tissue sections. *α*-SMA IF indicates activated MFs in the heart. N=100-115 cells were counted from multiple sections from murine hearts (n=6). (G) *Fam114a1* mRNA expression is induced in the hearts from mice with MI surgery at infarct zone (IZ), border zone (BZ), and remote zone (RZ) compared to the hearts with Sham surgery (20 days) (n=6). Data were presented as mean±SEM. Comparisons of means between two groups were performed by unpaired two-tailed Mann Whitney test for data not normally distributed for A, E, F, and unpaired Student *t* test for data normally distributed for B-D. Comparisons of means between >2 groups were performed by one-way ANOVA with Tukey’s multiple comparison test for G.

Immunofluorescence (IF) studies revealed that FAM114A1 protein expression was dramatically increased and strongly co-localized with *α*-SMA-positive cells (probably activated MFs), but not co- localized with cardiomyocytes (CMs) in human failing hearts compared to non-failure donor hearts (Figure 1E and S1A). We confirmed that severe fibrosis existed in human failing hearts as indicated by high collagen deposition compared to non-failure hearts (Figure S1B). To investigate whether FAM114A1 expression is associated with cardiac fibrosis during HF, Pearson correlation coefficient analysis was performed. The expression of *FAM114A1* mRNA showed a strong positive correlation with the expression level of collagen mRNAs such as COL1A1 (R=0.743) and COL3A1 (R=0.861) but not with that of ANP and BNP (Figure S1C). An independent dataset of human samples also showed increased expression of *FAM114A1* mRNA in the failing hearts of idiopathic dilated cardiomyopathy (IDCM) and ischemic cardiomyopathy (ICM) patients compared to healthy subjects (Figure S1D). Consistently, FAM114A1 protein was co-localized with *α*-SMA in Ang II-treated murine hearts, suggesting high level expression of FAM114A1 in activated MFs (Figure 1F). Besides, *FAM114A1* mRNA was highly induced in the ischemic myocardium in the mouse after MI surgery (Figure 1G). Intriguingly, the remote area showed the highest expression of *Fam114a1* mRNA. Moreover, FAM114A1 protein was significantly induced in the mouse ischemic myocardium (Figure S1E). Altogether, these findings indicate a strong positive correlation between FAM114A1 expression and cardiac fibroblast activation in both human and mouse heart failure.

### Fam114a1 deficiency in mice reduces cardiac fibrosis and protects cardiac function under pathogenic stresses

To determine the role of FAM114A1 in cardiac pathogenesis, we obtained *Fam114a1* global knockout (KO) mice created by the CRISPR-Cas9 technology (Figure S2A, B). The homozygous *Fam114a1*^−/−^ mice were viable, fertile, and normal in body weight and behavior. *Fam114a1* mRNA with a premature stop codon was detected by probes spanning exons 1-2 and 5-6 but not exons 2-3, suggesting a successful exon 3 deletion at both DNA and mRNA levels (Figure S2C). FAM114A1 protein expression was totally abolished in the heart of *Fam114a1*^−/−^ mice (Figure S2D). Thus, FAM114A1 is a non-essential gene and loss of both alleles does not cause developmental defects in mice at baseline.

Given the induction of FAM114A1 expression in human and mouse failing hearts, we ought to determine the pathophysiological effects of FAM114A1 deletion on the murine heart under cardiac remodeling *in vivo*. We first used Ang II to treat mice via osmotic minipump implantation (1.4 mg/Kg/day) for 4 weeks. The hearts of WT and *Fam114a1*^−/−^ mice were comparable in size at baseline (Figure S2E), as indicated by HW/TL (heart weight/tibia length) and HW/BW (heart weight/body weight) ratios (Figure 2A and S2F) as well as wheat germ agglutinin (WGA) staining (Figure 2B). Picrosirius red staining showed that *Fam114a1*^−/−^ mice had much less cardiac fibrosis than WT mice after Ang II treatment, suggesting that *Fam114a1*^−/−^ mice are more resistant to cardiac remodeling in response to Ang II stimulation than WT mice (Figure 2C). Baseline expression of multiple fibrosis marker genes (Col1a1, Col3a1, Ctgf, and Fn1) was not significantly altered in *Fam114a1*^−/−^ mice at baseline (Figure 2D). However, the expression of these genes was significantly lower in *Fam114a1*^−/−^ versus WT mice after Ang II treatment (Figure 2D). Also, expression of cardiac hypertrophy marker genes (Myh6, Myh7) was partially normalized in *Fam114a1*^−/−^ mouse hearts compared to those of WT mice upon Ang II treatment (Figure 2D). Echocardiographic examinations showed a significant decrease in the left ventricular ejection fraction (EF) and fractional shortening (FS) in Ang II-treated WT mice compared with vehicle-treated mice (Figure 2E, Table 1). In contrast, cardiac function was markedly restored in *Fam114a1*^−/−^ mice compared to WT mice after Ang II treatment (Figure 2E, Table 1).

**Figure 2.**
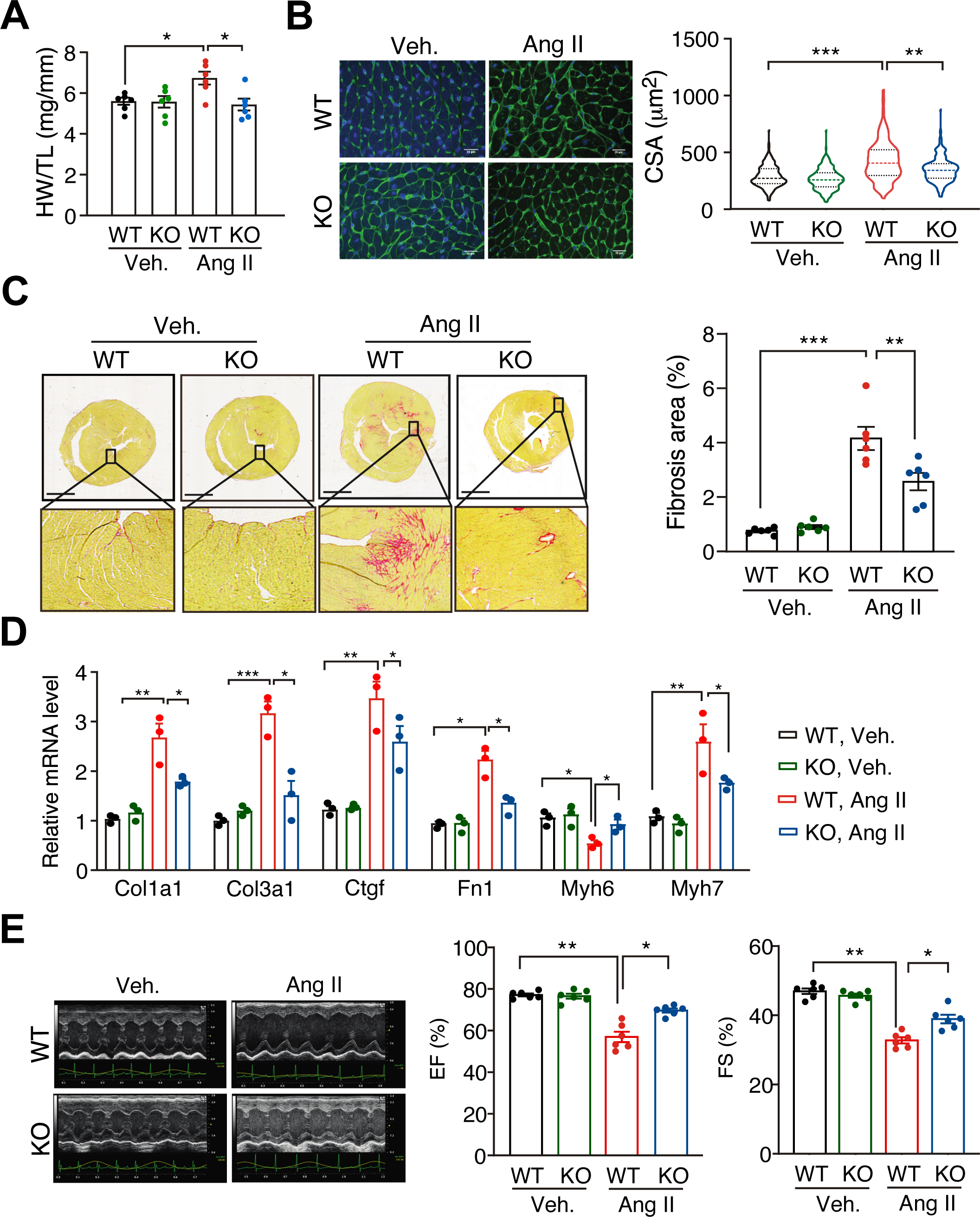
Deletion of FAM114A1 mitigates Ang II induced cardiac remodeling and heart failure in mice. (A) *Fam114a1*^−/−^ mice show reduced HW/TL (heart weight/tibia length) ratio after 4 weeks of Ang II infusion compared to WT mice. N=6 for each group. (B) WGA staining of heart tissue sections of WT and *Fam114a1^−/−^* mice under Ang II or vehicle treatment. Cardiac hypertrophy was measured based on quantified cross- sectional area of CMs. N=6 hearts per group with 250-270 CMs measured per heart. Scale bar: 20 μm. (C) Picrosirius red staining of heart tissue sections of WT and *Fam114a1^−/−^* mice under Ang II or vehicle treatment. Scale bar: 1 mm. N=6 for each group. (D) RT-qPCR measurement of cardiac fibrosis and hypertrophy marker gene expression in WT and *Fam114a1^−/−^* hearts after 4 weeks of Ang II versus vehicle infusion. *Rpl30* mRNA was used as a normalizer. N=3 for each group. (E) Representative echocardiographic images suggest improved cardiac function in Fam*114a1^−/−^* mice compared to WT mice after Ang II or vehicle infusion. Quantification of ejection fraction (EF) and fractional shortening (FS) was shown. N=6 for each group. Data were presented as mean±SEM. Statistical significance was confirmed by two-way ANOVA with Tukey’s multiple comparisons tests for A-E.

**Table 1.**
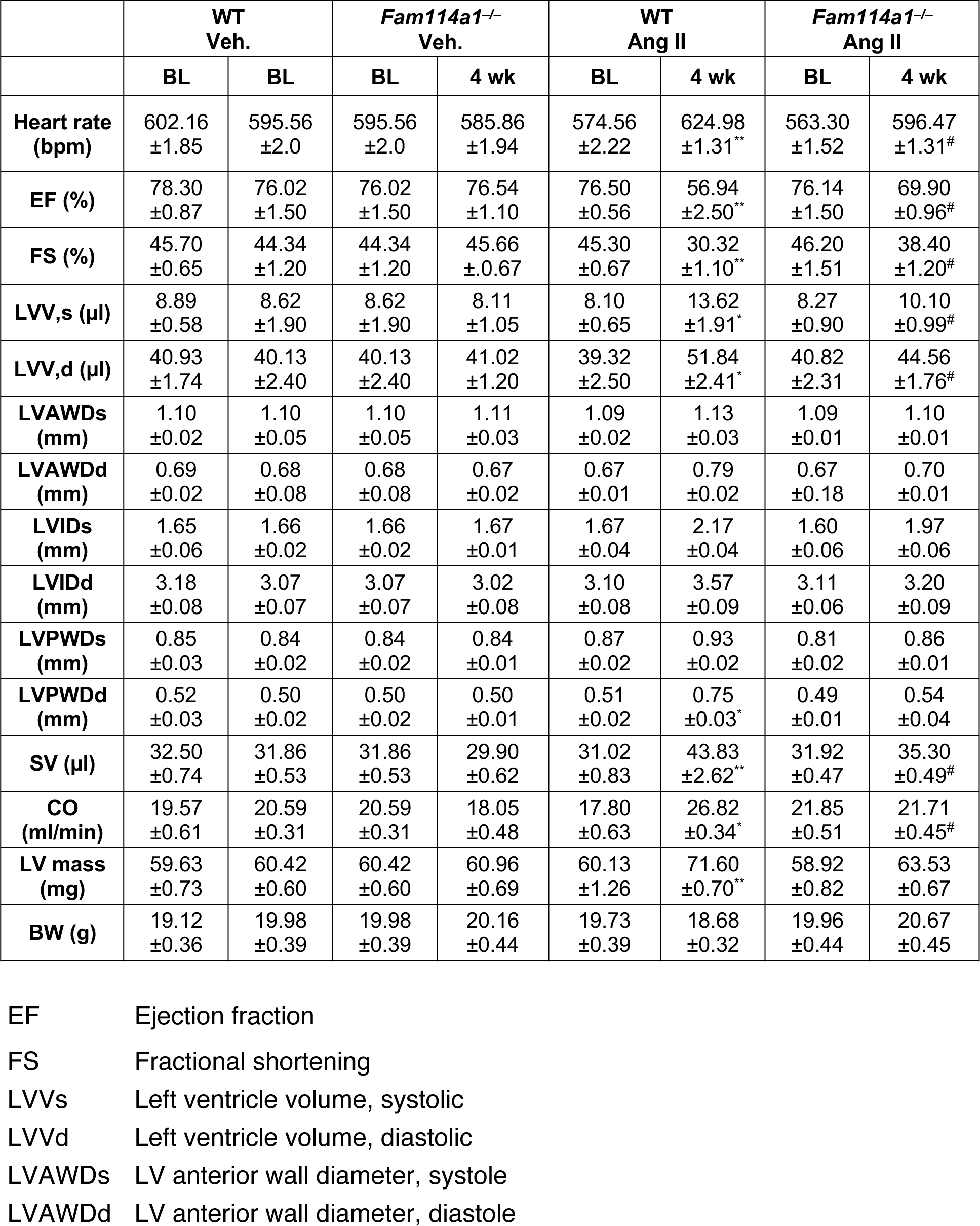
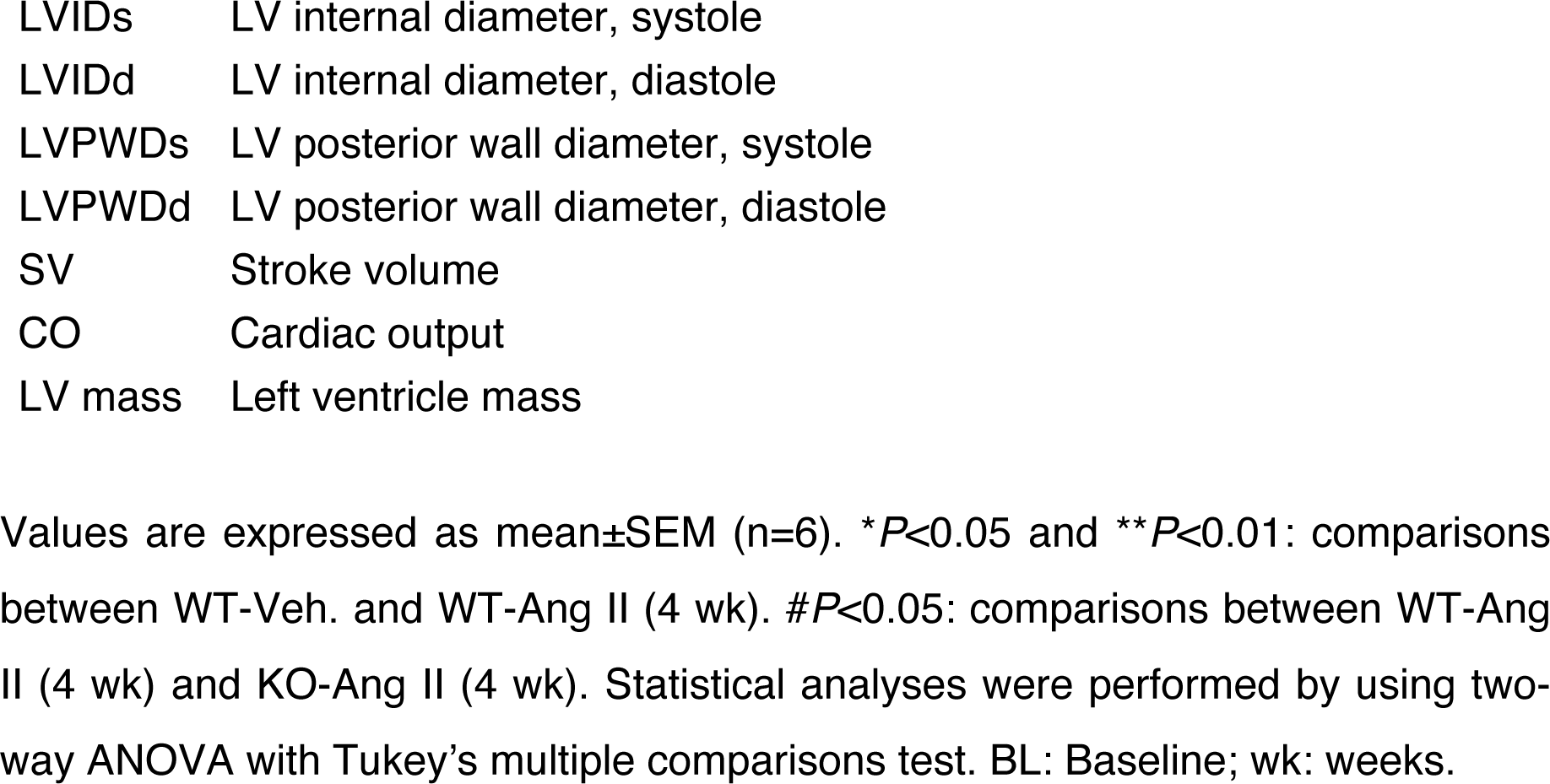
Echocardiographic analysis (M-mode short axis) of WT and *Fam114a1*^−/−^ mice after 4 weeks of Ang II infusion.

|  | WT Veh. |  | *Fam114a1*<sup>-/-</sup> Veh. |  | WT Ang II |  | *Fam114a1*<sup>-/-</sup> Ang II |  |
| --- | --- | --- | --- | --- | --- | --- | --- | --- |
|  | BL | BL | BL | 4 wk | BL | 4 wk | BL | 4 wk |
| **Heart rate (bpm)** | 602.16<br>±1.85 | 595.56<br>±2.0 | 595.56<br>±2.0 | 585.86<br>±1.94 | 574.56<br>±2.22 | 624.98<br>±1.31** | 563.30<br>±1.52 | 596.47<br>±1.31<sup>#</sup> |
| **EF (%)** | 78.30<br>±0.87 | 76.02<br>±1.50 | 76.02<br>±1.50 | 76.54<br>±1.10 | 76.50<br>±0.56 | 56.94<br>±2.50** | 76.14<br>±1.50 | 69.90<br>±0.96<sup>#</sup> |
| **FS (%)** | 45.70<br>±0.65 | 44.34<br>±1.20 | 44.34<br>±1.20 | 45.66<br>±0.67 | 45.30<br>±0.67 | 30.32<br>±1.10** | 46.20<br>±1.51 | 38.40<br>±1.20<sup>#</sup> |
| **LVV,s (μl)** | 8.89<br>±0.58 | 8.62<br>±1.90 | 8.62<br>±1.90 | 8.11<br>±1.05 | 8.10<br>±0.65 | 13.62<br>±1.91* | 8.27<br>±0.90 | 10.10<br>±0.99<sup>#</sup> |
| **LVV,d (μl)** | 40.93<br>±1.74 | 40.13<br>±2.40 | 40.13<br>±2.40 | 41.02<br>±1.20 | 39.32<br>±2.50 | 51.84<br>±2.41* | 40.82<br>±2.31 | 44.56<br>±1.76<sup>#</sup> |
| **LVAWDs (mm)** | 1.10<br>±0.02 | 1.10<br>±0.05 | 1.10<br>±0.05 | 1.11<br>±0.03 | 1.09<br>±0.02 | 1.13<br>±0.03 | 1.09<br>±0.01 | 1.10<br>±0.01 |
| **LVAWDd (mm)** | 0.69<br>±0.02 | 0.68<br>±0.08 | 0.68<br>±0.08 | 0.67<br>±0.02 | 0.67<br>±0.01 | 0.79<br>±0.02 | 0.67<br>±0.18 | 0.70<br>±0.01 |
| **LVIDs (mm)** | 1.65<br>±0.06 | 1.66<br>±0.02 | 1.66<br>±0.02 | 1.67<br>±0.01 | 1.67<br>±0.04 | 2.17<br>±0.04 | 1.60<br>±0.06 | 1.97<br>±0.06 |
| **LVIDd (mm)** | 3.18<br>±0.08 | 3.07<br>±0.07 | 3.07<br>±0.07 | 3.02<br>±0.08 | 3.10<br>±0.08 | 3.57<br>±0.09 | 3.11<br>±0.06 | 3.20<br>±0.09 |
| **LVPWDs (mm)** | 0.85<br>±0.03 | 0.84<br>±0.02 | 0.84<br>±0.02 | 0.84<br>±0.01 | 0.87<br>±0.02 | 0.93<br>±0.02 | 0.81<br>±0.02 | 0.86<br>±0.01 |
| **LVPWDd (mm)** | 0.52<br>±0.03 | 0.50<br>±0.02 | 0.50<br>±0.02 | 0.50<br>±0.01 | 0.51<br>±0.02 | 0.75<br>±0.03* | 0.49<br>±0.01 | 0.54<br>±0.04 |
| **SV (μl)** | 32.50<br>±0.74 | 31.86<br>±0.53 | 31.86<br>±0.53 | 29.90<br>±0.62 | 31.02<br>±0.83 | 43.83<br>±2.62** | 31.92<br>±0.47 | 35.30<br>±0.49<sup>#</sup> |
| **CO (ml/min)** | 19.57<br>±0.61 | 20.59<br>±0.31 | 20.59<br>±0.31 | 18.05<br>±0.48 | 17.80<br>±0.63 | 26.82<br>±0.34* | 21.85<br>±0.51 | 21.71<br>±0.45<sup>#</sup> |
| **LV mass (mg)** | 59.63<br>±0.73 | 60.42<br>±0.60 | 60.42<br>±0.60 | 60.96<br>±0.69 | 60.13<br>±1.26 | 71.60<br>±0.70** | 58.92<br>±0.82 | 63.53<br>±0.67 |
| **BW (g)** | 19.12<br>±0.36 | 19.98<br>±0.39 | 19.98<br>±0.39 | 20.16<br>±0.44 | 19.73<br>±0.39 | 18.68<br>±0.32 | 19.96<br>±0.44 | 20.67<br>±0.45 |

|  |  |
| --- | --- |
| LVIDs | LV internal diameter, systole |
| LVIDd | LV internal diameter, diastole |
| LVPWDs | LV posterior wall diameter, systole |
| LVPWDd | LV posterior wall diameter, diastole |
| SV | Stroke volume |
| CO | Cardiac output |
| LV mass | Left ventricle mass |

Values are expressed as mean ± SEM (n=6). \* $P$ <0.05 and \*\* $P$ <0.01: comparisons between WT-Veh. and WT-Ang II (4 wk). # $P$ <0.05: comparisons between WT-Ang II (4 wk) and KO-Ang II (4 wk). Statistical analyses were performed by using two-way ANOVA with Tukey's multiple comparisons test. BL: Baseline; wk: weeks.

To further confirm the phenotype of *Fam114a1*^−/−^ mice under Ang II infusion, we subjected *Fam114a1*^−/−^ mice to a more physiologically relevant HF model of permanent left anterior descending (LAD) coronary artery ligation-mediated MI. Ischemia-induced cardiac hypertrophy was significantly blunted in *Fam114a1*^−/−^ mice (Figure 3A and 3B). The fibrosis and infarct areas were markedly reduced in *Fam114a1*^−/−^ mice (Figure 3C and S3A, B). Consistently, fibrosis marker gene expression was reduced in *Fam114a1*^−/−^ mice, including *Col1a1* and *Col3a1* (Figure S3C). During MI, CFs start proliferation while CMs undergo apoptosis and cell death upon the cardiac injury. Therefore, we examined the cellular phenotypes of CFs in the *Fam114a1* KO mice. Less CF cell proliferation and CM cell death were observed in *Fam114a1*^−/−^ mice compared to WT mice after MI surgery (Figure 3D, E, and S3D). The cardiac function in *Fam114a1*^−/−^ mice was significantly improved compared to WT mice 14 days post-MI surgery indicated by partially recovered EF and FS as well as reduced left ventricle (LV) volumes (Figure 3F, S3E, and Table 2). In summary, *Fam114a1*^−/−^ mice exhibited reduced cardiac fibrosis, myocyte hypertrophy, and improved cardiac function in HF models under neurohumoral and ischemic stress conditions.

**Figure 3.**
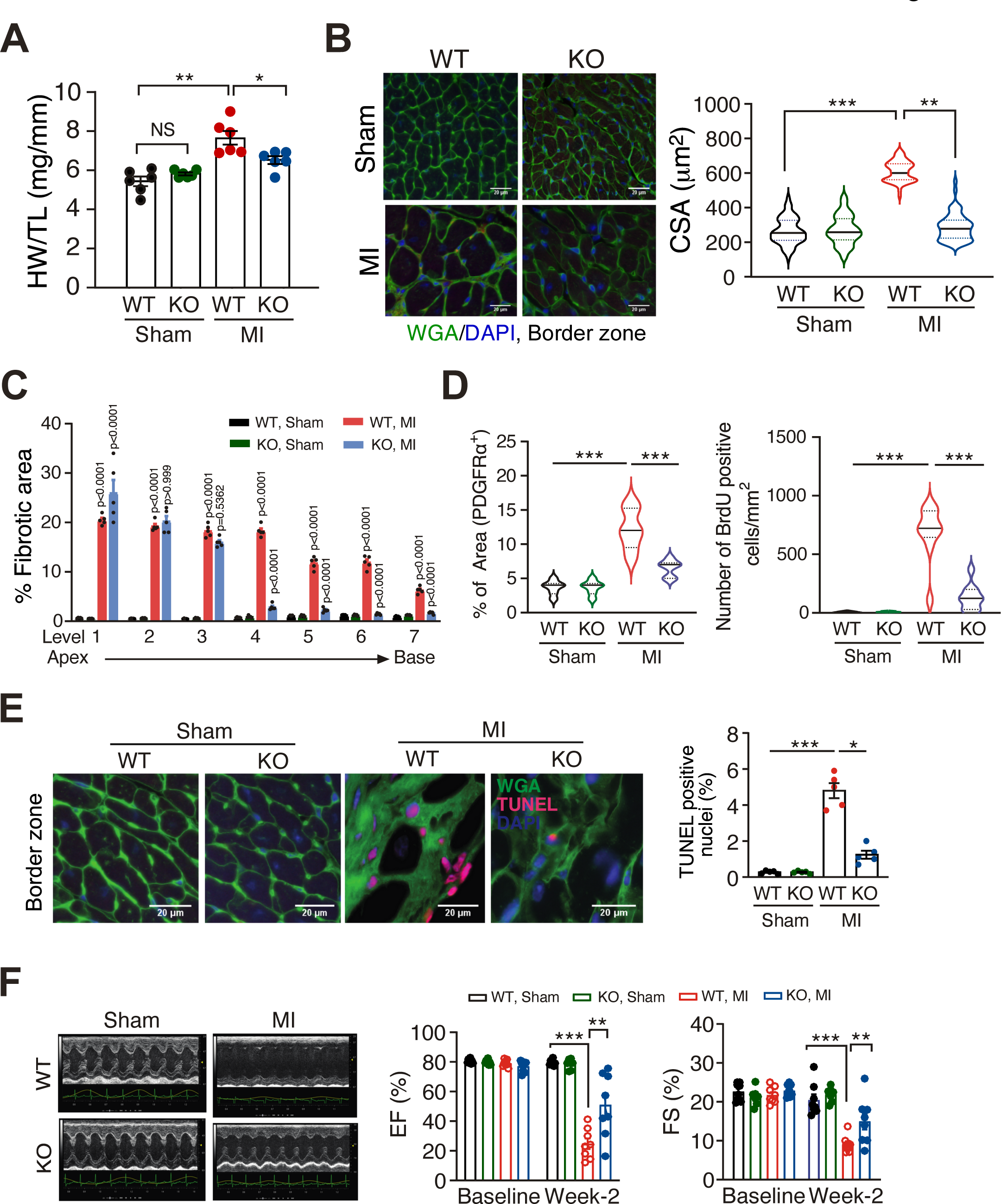
Deletion of FAM114A1 reduces cardiac remodeling and improves cardiac function after MI surgery. (A) *Fam114a1*^−/−^ mice show reduced HW/TL ratio after 20 days post MI surgery compared to WT mice. N=6 for each group. (B) WGA staining of heart tissue sections of WT and *Fam114a1^−/−^* mice after 20 days post MI surgery. N=6 for each group with 100-150 CMs in the border zone measured per heart. Scale bar: 20 μm. (C) Quantification of Picrosirius red staining of heart tissue sections of WT and *Fam114a1^−/−^* mice after 20 days post MI surgery. N=5 for each group. (D) Quantification of PDGFR*α* protein expression (% of the area) and number of BrdU positive cells in heart tissue sections of WT and *Fam114a1^−/−^* mice after 20 days post MI or Sham surgery. N=4 for each group and ∼100-150 positive cells were counted. (E) TUNEL assay for heart tissue sections (border zone) of WT and *Fam114a1*^−/−^ mice after 20 days post MI or sham surgery. Scale bar: 20 μm. N=5 for each group. (F) Representative echocardiographic images of WT and *Fam114a1*^−/−^ mice at baseline and after 2 weeks post MI or sham surgery. N=8 for each group. Quantification of EF and FS was shown. Data were represented as mean±SEM. Statistical significance was confirmed by two-way ANOVA with Tukey’s multiple comparisons tests for A-F.

**Table 2:**
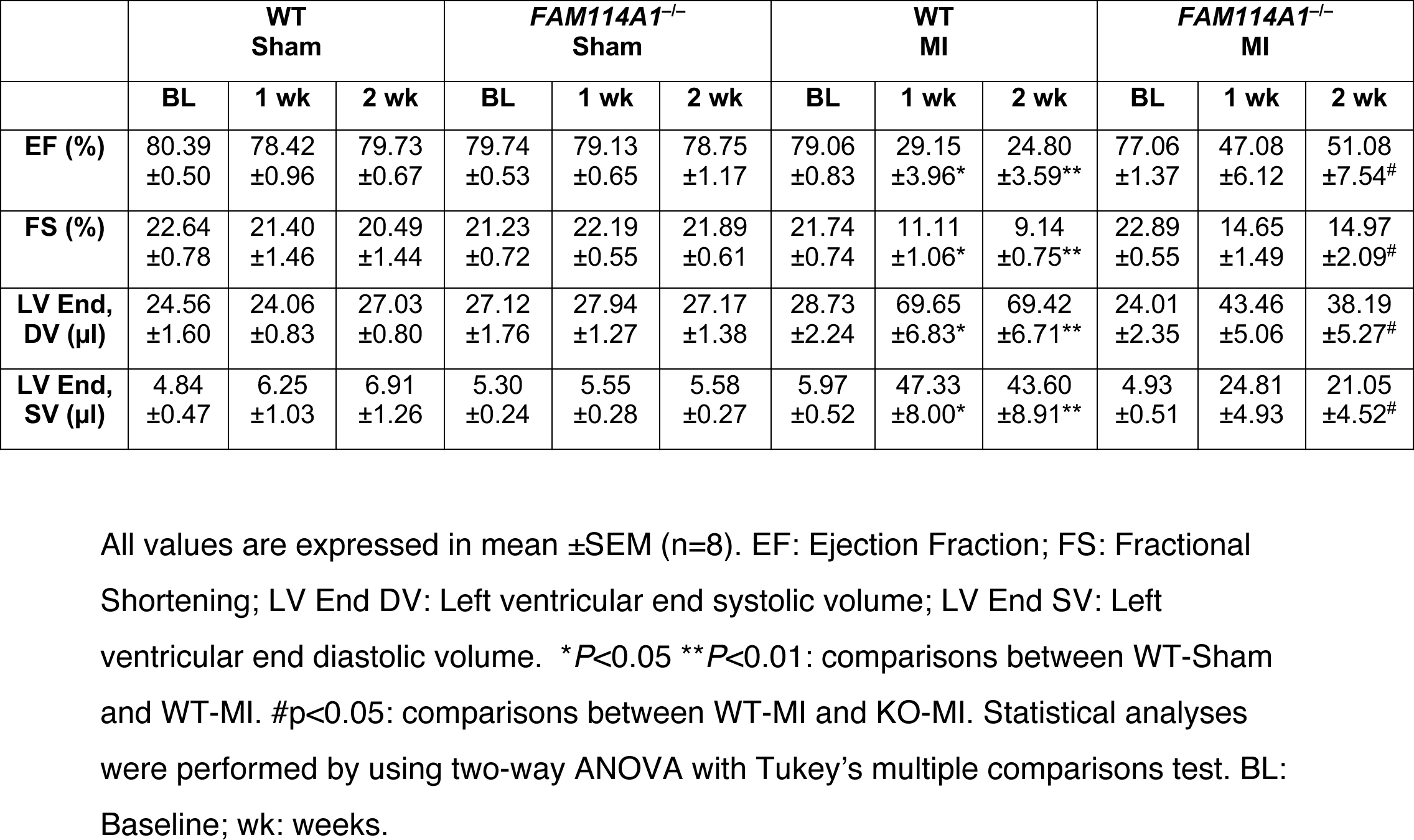
Echocardiographic analysis (B-mode long axis) of WT and *Fam114a1*^−/−^ after permanent LAD ligation.

### Fam114a1 deficiency in mice reduces cardiac inflammation and oxidative stress under pathogenic stresses

One major downstream effect of Ang II stimulation and MI is increase in inflammatory and oxidative responses and consequent injury to the heart. Thus, we examined whether FAM114A1 deletion influences inflammation and superoxide production in Ang II infused mouse hearts. We found that the mRNA expression of inflammatory genes (e.g., *Il-6*, Tnf*α*, Cxcl1, and Arg1) was reduced in *Fam114a1*^−/−^ hearts compared to WT hearts (Figure 4A). Cytokine array analyses in mouse serum samples showed that loss of FAM114A1 reduced circulating CXCL9 and CXCL10 levels (Figure S4A). Also, we observed less infiltration of CD45 positive leukocyte cells in the myocardium of *Fam114a1*^−/−^ mice compared to that of WT mice upon Ang II infusion, suggesting reduced inflammatory responses in the KO hearts (Figure 4B). Similarly, less immune cell infiltration was observed in both infarct and remote areas of MI hearts of *Fam114a1*^−/−^ mice compared to WT mice; while no difference was seen under sham conditions between the WT and *Fam114a1*^−/−^ mice (Figure S4B, C). Reactive oxygen species (ROS) are one of the primary triggers of inflammatory response in the heart after cardiac injury. Dihydroergotamine (DHE) staining showed that ROS production was remarkably induced in cultured WT CFs under Ang II treatment and significantly abolished in *Fam114a1*^−/−^ CFs (Figure 4C). One major source of ROS is produced by nicotinamide adenine dinucleotide phosphate (NADPH) oxidases such as NOX2 and NOX4. We measured the expression of NADPH oxidases and found that *Nox2* and *Nox4* mRNA levels were significantly reduced in *Fam114a1*^−/−^ hearts compared to WT hearts after Ang II infusion (Figure 4D). Altogether, these results suggest that deletion of *Fam114a1* ameliorates Ang II and MI induced cardiac inflammation and oxidative stress in the genetic animal model.

**Figure 4.**
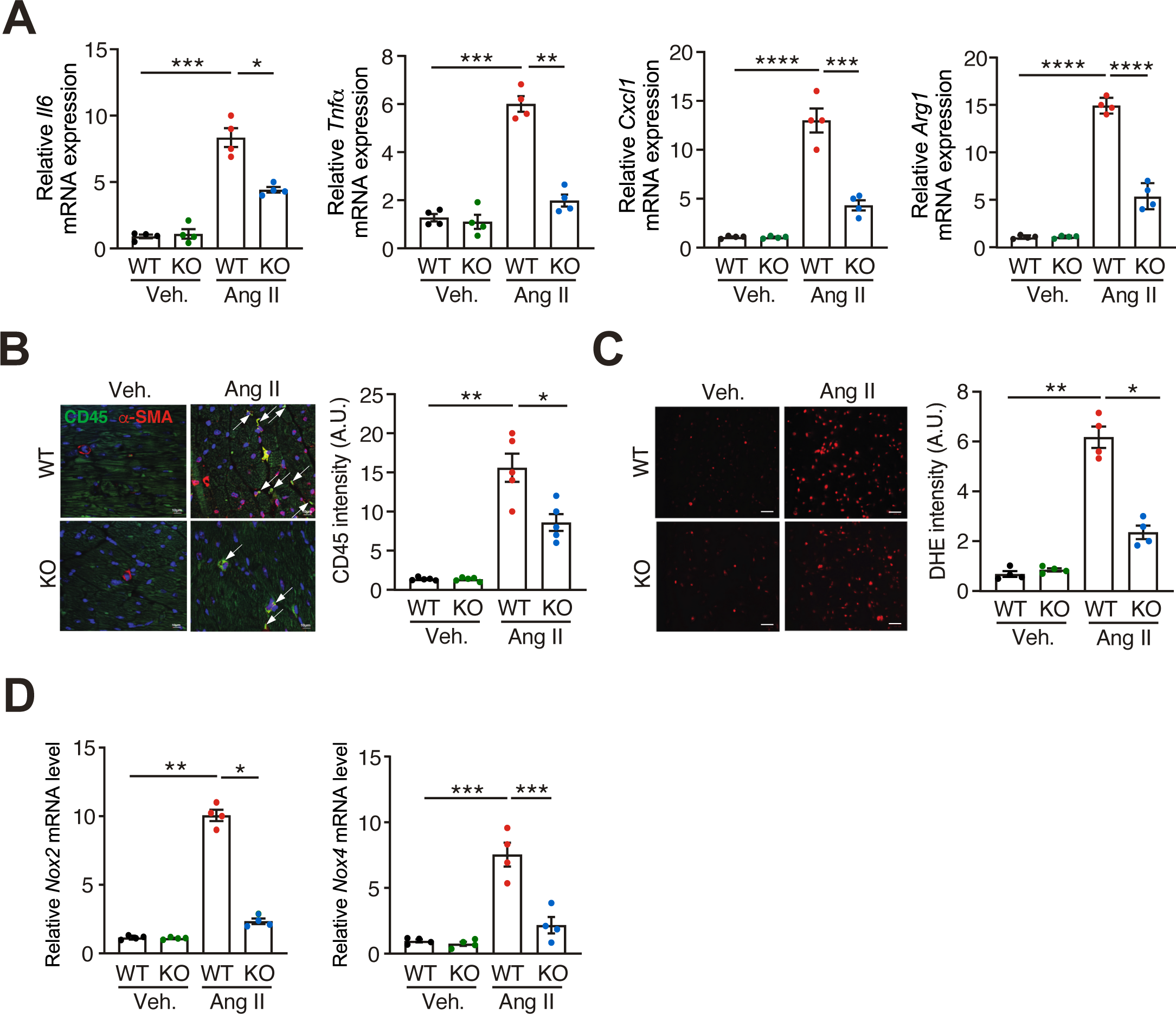
FAM114A1 deletion antagonizes Ang II induced inflammation and oxidative stress. (A) RT-qPCR measurement of mRNA expression levels of Il-6, Tnf-*α*, Cxcl1 and Arg1 in WT and *Fam114a1*^−/−^ mice. N=4 for each group. *Gapdh* mRNA was used as normalizer. (B) Representative IF images show CD45 (green) and *α*-SMA (red) in heart tissue sections of WT and *Fam114a1*^−/−^ mice after 4 weeks of Ang II infusion. Scale bar: 10 μm. N=5 for each group. (C) DHE staining of cultured CF cells isolated from WT and *Fam114a1*^−/−^ mice. Scale bar: 100 μm. N=4 for each group. (D) RT-qPCR measurement of mRNA expression level of NADPH oxidase genes (*Nox2* and *Nox4*). *Gapdh* mRNA was used as a normalizer. N=4 for each group. Data were represented as mean±SEM. Statistical significance was confirmed by two-way ANOVA with Tukey’s multiple comparisons tests for A-D.

### Loss of FAM114A1 in cardiac fibroblasts attenuates response to Ang II signaling transduction and inhibits fibroblast proliferation and activation

To determine the function of FAM114A1 in specific cardiac cell types, we first surveyed the Genevestigator database and found that *FAM114A1* mRNA was dominantly expressed in multiple human fibroblast cells from different organs (e.g., skin, liver, heart) and modestly expressed in immune cell types (e.g., T cells, monocytes) (Figure S5A, upper panel). We also examined the single cell RNA-Seq data from the Human Protein Atlas database and confirmed that *FAM114A1* mRNA was highly expressed in CFs but lowly expressed in CMs (Figure S5A, bottom panel). We further confirmed that Fam114a1 mRNA and protein were dominantly expressed in isolated primary mouse CFs and barely detected in primary mouse CMs (Figure 5A, B). This observation indicates that FAM114A1 may primarily regulate gene expression and cellular function in CFs. Thus, we isolated primary mouse CFs for *in vitro* Ang II treatment. Ang II significantly induced Fam114a1 mRNA and protein expression in cultured mouse CFs (Figure 5C). Expression of CF proliferation marker genes Ccna2 and Ccne1 and myofibroblast activation marker genes Acta2 and Col1a1 were significantly reduced in the CFs from *Fam114a1*^−/−^ hearts compared to those of WT hearts upon Ang II treatment (Figure 5D and S5B). Consequently, *α*-SMA and COL1A1 protein expression was markedly reduced in the CFs from *Fam114a1*^−/−^ hearts compared to those from WT hearts after Ang II treatment (Figure 5E, F). Moreover, the cell migration ability of *Fam114a1*^−/−^ CFs was weaker than that of WT CFs at baseline and upon Ang II treatment (Figure 5G). Taken together, our results indicate that FAM114A1 is dominantly expressed in CFs, inducible upon Ang II stimulation, and is required for efficient CF cell proliferation, activation and migration.

**Figure 5.**
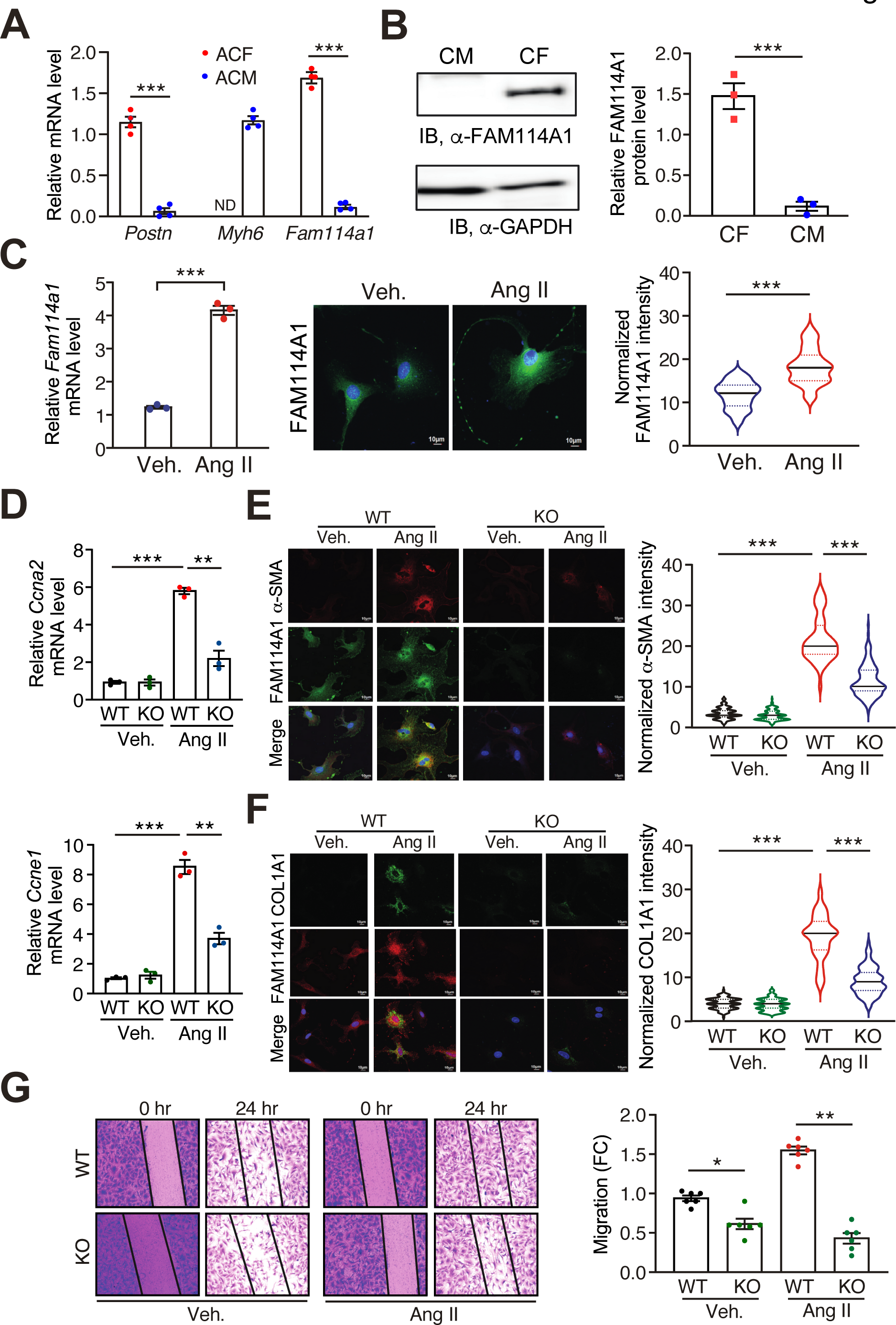
FAM114A1 deletion abolishes Ang II induced cardiac fibroblast proliferation, myofibroblast activation, and migration. (A) *Fam114a1* mRNA expression in primary mouse CFs and CMs. *Gapdh* mRNA was used as normalizer. N=4 for each group. (B) Endogenous FAM114A1 protein expression in primary mouse CFs and CMs detected by Western blot. GAPDH protein is used as a loading control. N=3 for each group. (C) Ang II treatment induces FAM114A1 mRNA and protein expression in primary mouse CFs. N=3 for RT-qPCR and n=100-120 cells per treatment for IF. 18S rRNA was used as normalizer for mRNA measurement. (D) RT-qPCR measurement of mitotic cyclin gene expression in CFs from WT and *Fam114a1*^−/−^ mice after Ang II treatment. N=3 for each group. 18S rRNA was used as normalizer. (E-F) Representative images of IF staining and quantification of normalized intensity of myofibroblast activation markers (*α*-SMA and COL1A1) in CFs of WT and *Fam114a1*^−/−^ mice after vehicle or Ang II treatment. N=150-200 cells were analyzed for each group. (G) Representative images of migrating CFs from WT and *Fam114a1^−/−^* mice after vehicle or Ang II treatment and quantification of scratch closure after 24 hrs was shown. N=6 for each group and each experiment is repeated 3 times. Data were represented as mean±SEM. Statistical significance was confirmed by two-tailed Student *t* test for A-C and two-way ANOVA with Tukey’s multiple comparisons test for D-G.

### FAM114A1 interacts with AGTRAP and regulates AGTRAP-AT1R expression in cardiac fibroblasts

Publicly available protein-protein interaction data have uncovered several potential interacting partners of FAM114A1, including CMTM5, RAB2B, SPG21, and AGTRAP among others (Figure S6A) (24). These candidate interacting proteins were discovered by yeast two hybrid screen, suggesting a direct physical interaction between the candidate protein and FAM114A1. Out of four tested candidate proteins, we validated the interaction between AGTRAP and FAM114A1 in mouse heart lysates and cultured primary mouse CFs by immunoprecipitation (IP) and Western blotting (Figure 6A and S6B). FLAG-tagged FAM114A1 recombinant protein was also detected to be associated with AGTRAP when overexpressed in NIH/3T3 fibroblast cells (Figure S6C). AGTRAP has been reported to bind AT1R and promote its internalization and degradation (21-23). Therefore, we ought to examine the role of FAM114A1 in regulation of the Ang II signaling via interacting with AGTRAP. At baseline, EGFP-tagged FAM114A1 was partially co-localized with Myc-tagged AGTRAP in vehicle-treated NIH/3T3 fibroblast cells (Figure 6B). Upon Ang II stimulation of fibroblast cells, expression level of EGFP-tagged FAM114A1 was enhanced possibly due to post-transcriptional regulation while Myc-tagged AGTRAP level was decreased in the cytoplasm (Figure 6B). Using primary mouse CF cell culture, we found that AGTRAP protein expression (but not mRNA) was increased in Ang II-treated *Fam114a1*^−/−^ CFs compared to WT CFs (Figure 6C and S6D). In contrast, AT1R protein expression was decreased in Ang II-treated *Fam114a1*^−/−^ CFs (Figure 6C). More importantly, we observed decreased AGTRAP and increased AT1R together with induced FAM114A1 protein expression in human failing hearts compared to non-failure donor hearts (Figure S6E). These observations in human samples are consistent with the data obtained in mouse heart failure models. Collectively, these results suggest that FAM114A1 interacts with AGTRAP in CFs and loss of FAM114A1 increases AGTRAP protein expression and may promote AT1R protein turnover.

**Figure 6.**
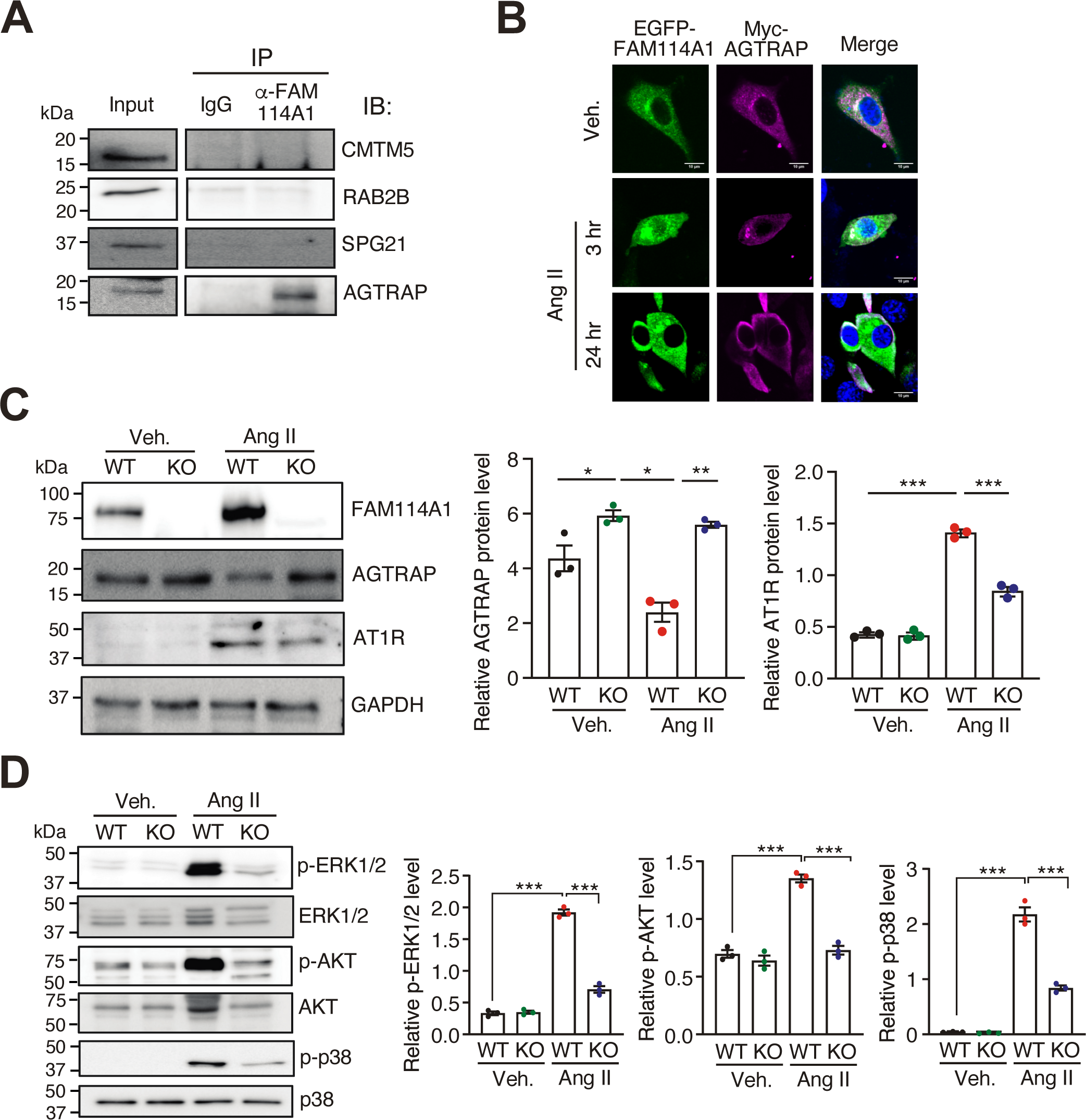
FAM114A1 depletion stabilizes AGTRAP, enhances AT1R degradation, and influences Ang II signaling in cardiac fibroblasts. (A) Immunoprecipitation and immunoblot confirm the interaction of FAM114A1 with AGTRAP in the mouse heart tissue lysates. (B) IF detection of overexpressed EGFP-tagged FAM114A1, Myc-tagged AGTRAP, and FLAG-tagged AT1R in mouse NIH/3T3 fibroblast cells. Protein expression changes were observed after 3 hrs and 24 hrs post Ang II or vehicle treatment. Scale bar: 10 μm. (C) Western blot measurement and quantification of FAM114A1, AGTRAP, and AT1R in vehicle or Ang II treated primary CF cells isolated from WT and *Fam114a1^−/−^* mice. (D) Western blot measurement and quantification of p-ERK1/2, ERK1/2, p-AKT, AKT, p- p38 and p38 in Ang II or vehicle treated CFs (24 hrs) from hearts of WT and *Fam114A1^−/−^* mice (n=3). Data were represented as mean±SEM. Statistical significance was confirmed by two-way ANOVA with Tukey’s multiple comparisons test for C-D.

Since FAM114A1 regulates protein expression of AGTRAP and AT1R, we next examined the role of FAM114A1 in Ang II-induced signaling transduction in cultured primary CFs from WT and *Fam114a1*^−/−^ mice. Our immunoblot results reveal that in Ang II-treated *Fam114a1*^−/−^ CFs phosphorylation of ERK1/2, AKT, and p38 was significantly abolished compared to WT CFs (Figure 6D). These data suggest that the deletion of *Fam114a1* antagonizes the Ang II-induced activation of downstream effectors and hence reduces the collagen deposition.

### Transcriptome profiling reveals dysregulated gene pathways in Fam114a1 null cardiac fibroblasts

To further understand global gene expression changes in *Fam114a1* null CF cells, we performed RNA-Seq in isolated primary adult CFs from WT and *Fam114a1*^−/−^ mice. Multiple dysregulated genes were uncovered at the steady-state mRNA level in *Fam114a1*^−/−^ CFs (Figure 7A and 7B). *Fam114a1* mRNA was reduced by 4 folds probably due to a robust nonsense-mediated mRNA decay triggered by the presence of a premature termination codon introduced by CRISPR-Cas9- mediated gene editing (Figure S2A). Several genes showed more drastic reduction of mRNA expression compared to Fam114a1, such as Ifi208 (interferon activated gene 208), Adamts15 (ADAM metallopeptidase with thrombospondin type 1 motif 15), Serpinc1 (serpin family C member 1), among others (Table S1). David gene ontology analyses suggest that the downregulated genes are mainly involved in two pathways, including basement membrane and extracellular matrix (Figure 7C). The decrease of a select of genes was confirmed by quantitative RT-PCR (Figure 7D).

**Figure 7.**
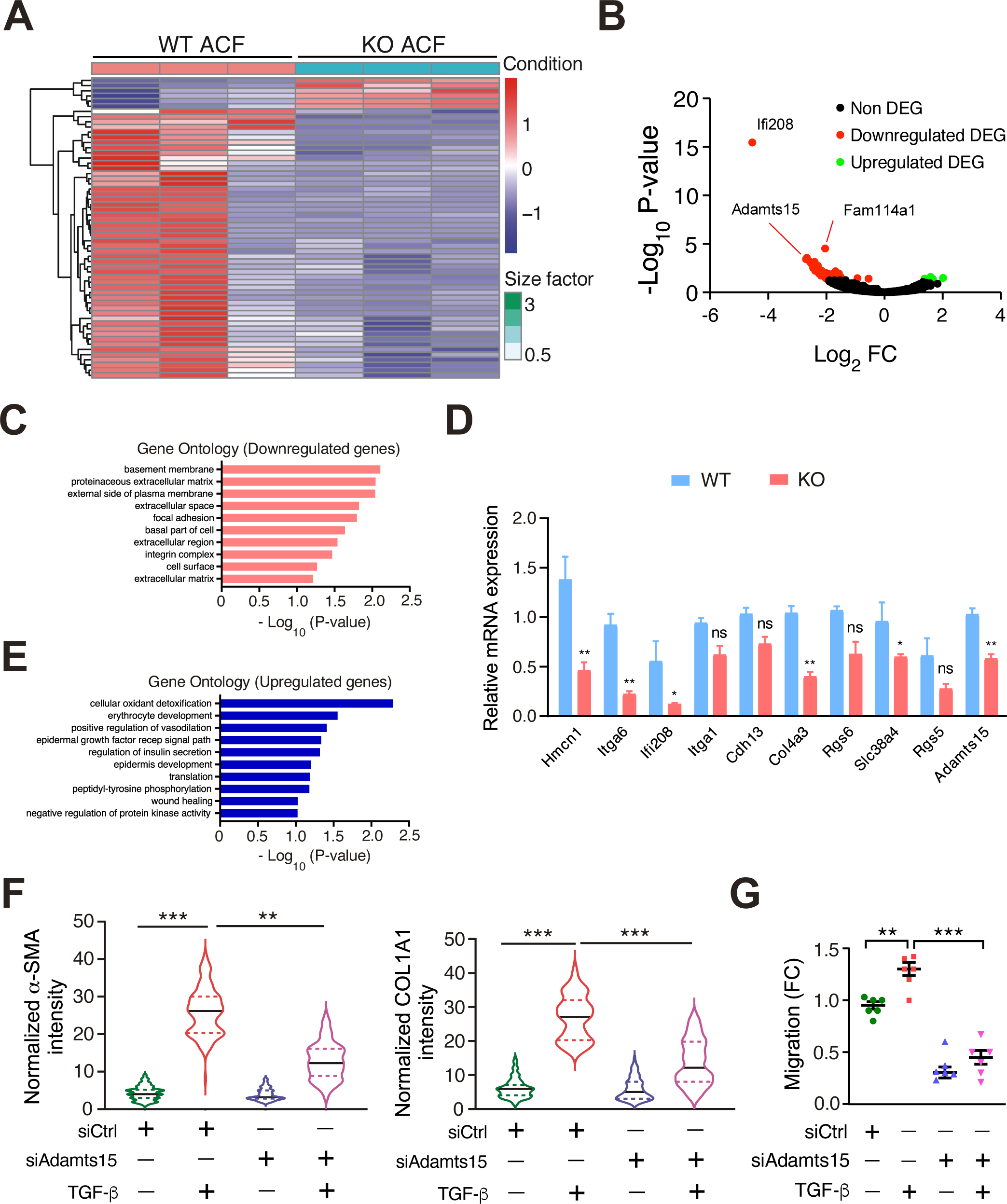
ADAMTS15, a differentially regulated genes in *Fam114a1* null cardiac fibroblasts, is required for CF-to-myofibroblast activation. (A) Heatmap of gene expression in CFs isolated from WT and *Fam114a1*^−/−^ mice at baseline analyzed by RNA-Seq. P60 male mice, n=3 per group, *P*adj<0.05. (B) Volcano plot of differentially expressed genes in CFs isolated from WT and *Fam114a1*^−/−^ mice at baseline analyzed by RNA-Seq. (C) Gene Ontology analysis of enriched pathways of downregulated genes in RNA-Seq. The top pathways are listed with enriched gene sets. (D) RT-qPCR validation of multiple downregulated genes in *Fam114a1*^−/−^ mice derived CFs. 18S rRNA was used as a normalizer (n=3). (E) Gene Ontology analysis of enriched pathways of upregulated genes in RNA-Seq. (F) IF quantification of COL1A1 and *α*-SMA protein expression after knockdown of Adamts15 in TGF-*β*-treated primary mouse CFs. N=100-120 cells from three biological replicates were analyzed. (G) Knockdown of Adamts15 reduced primary mouse CF cell migration indicated by scratch assays (n=6). Data were represented as mean±SEM. Statistical significance was confirmed by unpaired Student *t* test for D and one-way ANOVA with Tukey’s multiple comparisons test for F, G.

On the other hand, we also observed increased gene expression of cellular oxidant detoxification and positive regulation of vasodilation among others (Figure 7E), which implies a compensatory response after FAM114A1 depletion *in vivo*.

### FAM114A1 downstream effector ADAMTS15 is required for cardiac fibroblast activation and migration

Findings of FAM114A1-mediated regulation of CF activation prompted us to further identify critical downstream effectors involved in this pathological process. Among the validated downregulated genes in *Fam114a1* null CFs, Adamts15 is among the most downregulated genes with high baseline mRNA expression. Furthermore, it has been identified as a potential modulator of hepatic fibrosis (33). Therefore, we ought to test whether ADAMTS15 plays a role in cardiac fibrosis. We found that *ADAMTS15* mRNA expression was increased in human failing hearts compared to non-failure donor hearts (Figure S7A) and positively correlated with *ACTA2* mRNA expression (Figure S7B). ADAMTS15 protein was colocalized with *α*-SMA positive CF cells in human failing hearts (Figure S7C) and mouse MI hearts (Figure S7D). siRNA-mediated knockdown of ADAMTS15 in mouse primary CFs significantly reduced protein expression of *α*-SMA and COL1A1 upon TGF-*β* treatment (Figure 7F and S7E). In addition, ADAMTS15 knockdown remarkably inhibited CF cell migration at baseline and after TGF-*β* stimulation (Figure 7G). Taken together, we discovered that FAM114A1 depletion in CFs reduces ECM gene expression and one critical downstream effector gene ADAMTS15 is required for TGF-*β*-triggered CF activation and migration.

## Discussion

In this study, we found a novel gene FAM114A1 that plays an important role in pathological cardiac remodeling and fibrosis. FAM114A1 is induced in human failing hearts as well as in HF mouse models of Ang II and MI. We used a global *Fam114a1* KO mouse model to demonstrate that deletion of FAM114A1 antagonizes pathological cardiac remodeling, fibrosis, and inflammation *in vivo*. We showed high expression of FAM114A1 in CFs but not in CMs. We have identified AGTRAP as a direct interacting partner of FAM114A1 in CFs. FAM114A1 downregulates the expression of AGTRAP and increases AT1R level, thereby enhancing Ang II signaling (Figure 8). Based on transcriptome profiling in *Fam114a1* null CFs, we identified multiple dysregulated genes that are involved in cardiac fibrosis, including a critical ECM protein ADAMTS15 that is required for TGF-*β*-triggered CF activation and migration. Our work indicates the importance of FAM114A1 in the pathogenesis of cardiac fibrosis and establishes FAM114A1 as a novel therapeutic target for heart disease.

**Figure 8.**
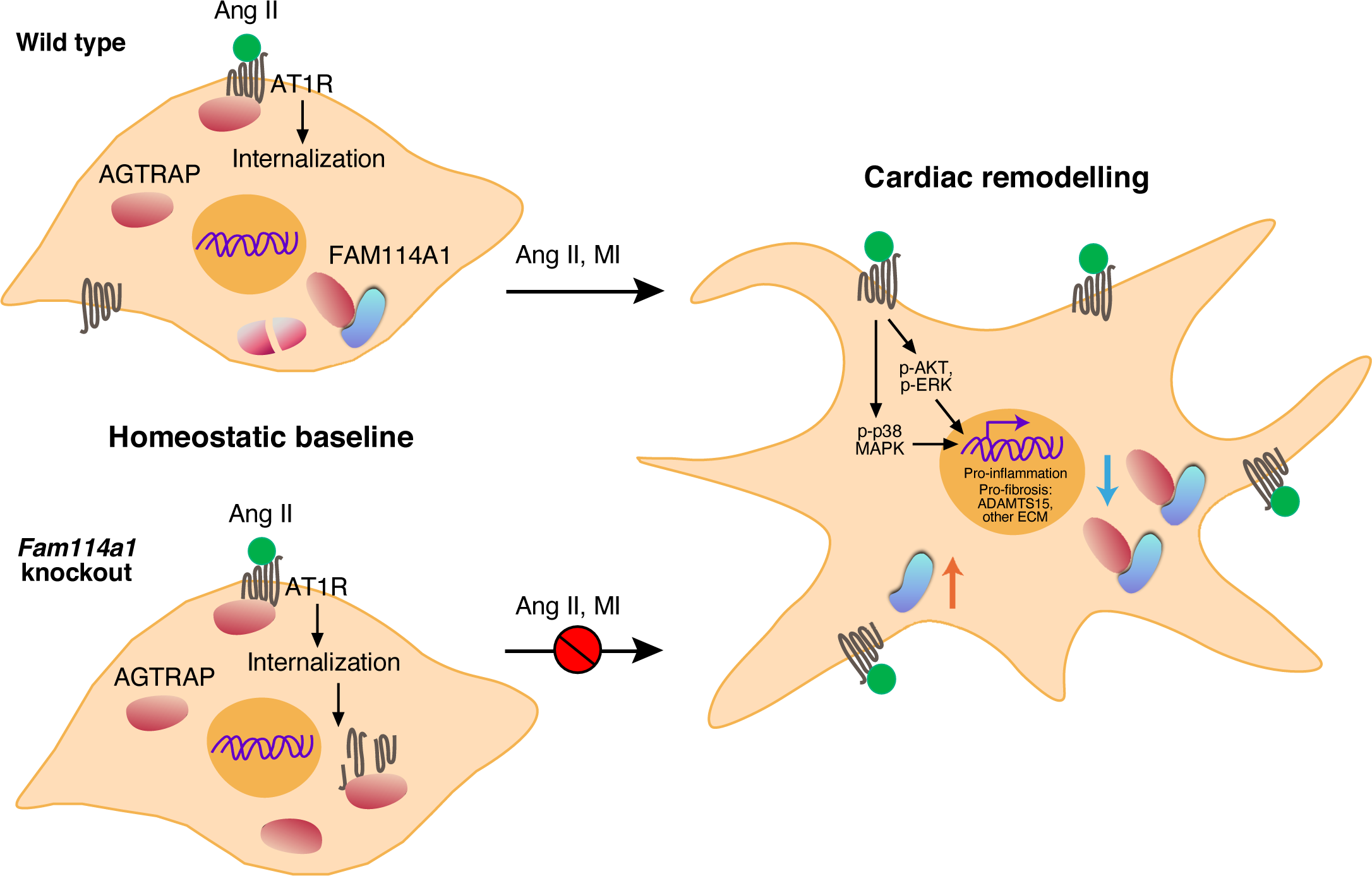
Schematic model of FAM114A1-mediated regulation of Ang II signaling and cardiac fibrosis.

To date, multiple signaling pathways have been reported to induce cardiac fibrosis, including TGF-*β* (34), ERK (35), signal transducer and activator of transcription 3 (STAT3) (36), SMAD2/3 (37), p38 MAPK (38), and EPRS (39). Since all these signaling pathways are downstream of Ang II signaling transduction and prominently affected by the activation of the angiotensin receptor (34), therapeutic strategies targeting molecules associated with Ang II signal transduction are promising to treat cardiac fibrosis and prevent arrhythmia. Our studies have two major conceptual advances in treating cardiac fibrosis and HF via abolishing FAM114A1-mediated amplification of Ang II signaling. First, this work is a pioneering study of the function of FAM114A1 and FAM114A1- AGTRAP axis that modulates cardiac fibrosis, inflammation, and cardiac remodeling. FAM114A1 is a novel regulatory factor of angiotensin signaling in cardiac fibrosis through modulating AGTRAP protein expression in CFs. Under cardiac stresses, increased FAM114A1 promotes cardiac fibrosis and pathological remodeling through the FAM114A1-AGTRAP-AT1R axis in CFs (Figure 8). Second, we show that FAM114A1 is induced upon multiple pathological cardiac stresses. Using RNA-Seq in isolated mouse primary CFs, we discover that FAM114A1-regulated genes and pathways are mainly involved in extracellular matrix and space (e.g., Adamts15, Col4a3, Hmcn1, Serpinc1, etc.). We have uncovered novel genes such as Adamts15 that is involved in CF activation. Taken together, these novel findings offer a strong rationale for FAM114A1 to be considered as a drug target in treating cardiac fibrosis and HF. At the mechanistic level, we will further characterize the molecular basis of FAM114A1-mediated downregulation of AGTRAP at post-transcriptional level. Post-translational modifications of FAM114A1 or AGTRAP and potential involvement of E3 ubiquitin ligases upon Ang II stimulation in CFs will need to be investigated in the future.

*FAM114A1* is a host gene for miR-574 and shares the common promoter with embedded intronic miR-574 (31, 32). Both the host gene *FAM114A1* and miR-574 expression is correlated (32). Thus, it is expected that *FAM114A1* mRNA is induced in human and mouse MI and even aged murine hearts together with miR-574 (28-32). We have confirmed that FAM114A1 mRNA and protein expression is induced in human and mouse failing hearts compared to healthy hearts, especially in the activated MFs (Figure 1 and S1). We previously observed induced miR-574-3p in activated CFs in WT mice upon the transverse aortic constriction (TAC) surgery (31). These results suggest that the host gene FAM114A1 and embedded intronic miR-574-3p are coordinated in their expression. Intriguingly, miR-574-3p plays a role in antagonizing TGF-*β*-induced CF activation in primary cell culture as well as in TAC mouse model via nanoparticle delivery (31). Here, we show that the host gene FAM114A1 contributes to enhanced Ang II signaling and pro-fibrotic response, implying an antagonistic function relationship between the host gene and embedded miRNA. In our *Fam114a1*^−/−^ hearts, we did not observe any significant change in the expression of miR-574- 3p or miR-574-5p (data not shown). This is because splicing of intron 1 or processing of miR-574 precursor occurs in the nucleus prior to the nonsense-mediated mRNA decay of the exon 3- removed *Fam114a1* mRNA in the cytoplasm. Therefore, the phenotypes we observed in *Fam114a1*^−/−^ mice are not caused by change of expression of miR-574.

FAM114A1 has multiple single nucleotide polymorphisms (SNPs) that are associated with human inflammatory diseases such as ankylosing spondylitis and host immune response to bacterial infection (40, 41). Importantly, human genome-wide association study databases (GWASATLAS) demonstrate that FAM114A1 has a genetic SNP mutation related to hypertension phenotype in human coronary artery disease (ranked as 89 out of 19057 genes, *P*=3.64e-9) (Figure S8A) (25). In another independent study, one FAM114A1 SNP (rs1873197; *P*=3.32e-06) was identified as a CAD associated loci with CAD diagnosis (e.g., MI, acute coronary syndrome, chronic stable angina, or coronary stenosis >50%) (26). Excitingly, the genetic association of the same FAM114A1 SNP (rs1873197) with MI was further confirmed by a recent large-scale genome-wide analysis (n=639,000 human MI subjects; effect allele frequency is 6.7743%; *P*=8.48e-05) (Figure S8B) (27). These GWAS data suggest potential functional connection of FAM114A1 with immune-related or vascular cell types. The Ang II infusion model involves multiple cardiac cell types in addition to CMs and CFs such as endothelial cells (ECs) and smooth muscle cells (SMCs) because Ang II signaling acts on both vascular and cardiac cell types. We used isolated mouse primary CFs from *Fam114a1* KO mice to provide the proof of cell type-specific autonomous effects in CFs (Figure 5). To further support this idea, we used the MI model which is less dependent on SMCs compared to the Ang II model. We also demonstrated similar anti-fibrotic effects of deletion of *Fam114a1* in MI mice. Obviously, FAM114A1 is modestly expressed in immune cells including monocyte, T-cells, and B-cells (Figure S5A), indicating that most likely minor effects of FAM114A1 in immune cell types may contribute to cardiac phenotypes if any. In the future, the function of FAM114A1 in other vascular cell types warrant further studies to fully understand its biological role.

Based on transcriptome-wide profiling of isolated *Fam114a1* null CF cells, we discover that one downstream effector of FAM114A1, ADAMTS15, plays an important role in regulating CF activation and migration. We also observed increased gene expression of cellular oxidant detoxification and positive regulation of vasodilation, suggesting a compensatory response that may contribute to the relief of oxidative stress and pressure overload during cardiac pathogenesis. Global *Fam114a1* knockout animals are normal in development and do not exhibit pathological changes. This indicates that FAM114A1 is a proper drug target for anti-fibrosis treatment without triggering deleterious side-effects when inhibited or depleted.

## Materials and Methods

Further information is available in *SI Materials and Methods*.

### Human Specimens

All human samples of frozen cardiac tissues, including 20 samples from explanted failing hearts (10 ischemic and 10 dilated cardiomyopathy hearts) and 8 samples from non-failing donor hearts, as well as paraffin section slides from DCM (n=5), ISHF (n=5) or non-failing hearts (n=7) were acquired from the Cleveland Clinic. All surgical procedures and tissue harvesting were performed following the Cleveland Clinic procedure and guideline. This study was approved by Material Transfer Agreement between the URMC and the Cleveland Clinic. In Figure 1A, total RNA samples from two failing hearts showed degradation during the quality control process and were excluded from RT-qPCR analysis. All human samples were picked randomly based on the presence or absence of heart failure by our collaborator, Dr. Waihong Wilson Tang at Cleveland Clinic. We are blinded from any clinical data. The identity numbers (IDs) of the frozen cardiac tissues and histology tissues are listed below.

### Mice

The global *Fam114a1* (family with sequence similarity 114, member A1) KO mice were created by CRISPR technology in International Mouse Phenotyping Consortium (IMPC). The alteration caused the deletion of 314 bp, which results in the deletion of exon 3, amino acid change after residue 116, and early termination 3 amino acids later. For experiments with *Fam114a1*^−/−^ mice, control mice of the same age and gender from littermates or sibling mating were used. All animal procedures were performed in accordance with the National Institutes of Health (NIH) and the University of Rochester Institutional guidelines. Two different disease models of heart failure were used: (1) Ang II infusion model (1.4 μg/g/day). (2) Permanent LAD ligation model (MI). Both Ang II and MI models were performed by the Microsurgical Core in a double-blinded manner.

### Echocardiography

For Ang II infusion mouse model, M-mode short axis echocardiographic image collection was performed using a Vevo2100 echocardiography machine (VisualSonics, Toronto, Canada) and a linear-array 40 MHz transducer (MS-550D). Heart rate was monitored during echocardiography measurement. Image capture was performed in mice under general isoflurane anesthesia with heart rate maintained at 500-550 beats/min. LV systolic and diastolic measurements were captured in M-mode from the parasternal short axis. Fraction shortening (FS) was assessed as follows: % FS = (end diastolic diameter - end systolic diameter) / (end diastolic diameter) x 100%. Left ventricular ejection fraction (EF) was measured and averaged in both the parasternal short axis (M-Mode) using the tracing of the end diastolic dimension (EDD) and end systolic dimension (ESD) in the parasternal long axis: % EF=(EDD-ESD)/EDD. Hearts were harvested at multiple endpoints depending on the study. In addition to EF and FS, left ventricular end-diastolic diameter (LVEDD), left ventricular end-systolic diameter (LVESD), wall thickness of left ventricular anterior (LVAWT) and posterior (LVPWT) were also assessed. For MI mouse model, B-mode long axis echocardiographic imaging was performed and left ventricle EF, FS, end systolic volume (LVESV), and end diastolic volume (LVEDV) were measured.

### Picrosirus Red staining

Paraffin-embedded heart tissue sections were deparaffinized and the sections were incubated with Picrosirius red reagent (Abcam) for 1 hr at RT. Slides were then washed with 1% acetic acid followed by 100% ethyl alcohol and mounted with mounting medium. Images were captured using the Prime Histo XE Slide Scanner (Carolina) and fibrosis area was measured by Image J software (NIH, USA).

### Cell culture and transfection

NIH/3T3 cells were cultured in DMEM supplemented with 10% bovine calf serum (VWR) and 1% penicillin/streptomycin (ThermoFisher). Primary CFs isolated from mouse hearts of both genders were cultured in DMEM supplemented with 10% FBS (ThermoFisher) and 1% penicillin/streptomycin. Primary cells were used at P0 for CF activation assays. We use the Polyjet to transfer the NIH 3T3 cells along with DNA plasmids. siRNA transfection (100 nM) in primary CFs was performed using lipofectamine 3000 following the manufacturer’s instructions.

### Cardiac fibroblast activation assay

Adult cardiac fibroblasts were isolated from ∼2-3 months old mice of WT and *FAM114A1*^−/−^ and placed in a 35 mm glass-bottom dishes. After 2 hrs, attached cells were washed with 1x PBS for 3 times, changed to fresh CF culturing medium (DMEM containing 10% FBS and 1% penicillin/streptomycin), and cultured at 37°C for 12 hrs (or overnight). Then, cells were treated with serum starvation for 12 hrs followed by stimulation with Ang II (1 μM) for 24 hrs. Immediately cells were fixed with 4% paraformaldehyde (PFA) for immunofluorescence staining and/or lysed in TRIzol reagent for RNA isolation and to detect the myofibroblast activation markers gene expression levels by RT-qPCR.

### Statistical Analysis

All quantitative data were presented as mean±SEM, and analyzed using the GraphPad Prism (9.0.0) software. For comparison between 2 groups, unpaired two-tailed Mann-Whitney test for not normally distributed data and unpaired two-tailed Student t-test for normally distributed data were performed. For multiple comparisons among ≥3 groups, Tukey’s one-way or two-way ANOVA with multiple comparisons test was performed. Statistical significance was assumed at a value of *P*<0.05.

## DATA AVAILABILITY

RNA-Seq data produced and used in this study were deposited on under accession number GSE165074 (https://www.ncbi.nlm.nih.gov/geo/query/acc.cgi?acc=GSE165074) in Gene Expression Omnibus (GEO) database and are scheduled to be released on Apr 19, 2021

## ACKNOWLEDGMENTS

We are grateful to Qiuqing Wang (Cleveland Clinic) and Virginia Aswad for biostatistical consulting and critical reading of the manuscript. We appreciate the technical assistance from Erika Flores Medina (Aab Cardiovascular Research Institute), Mohan Amy, and Deanne Mickelsen (Aab Cardiovascular Research Institute) in histology and surgical operations, respectively. RNA sequencing and primary data analysis were performed by Jason R Myers and Cameron Baker from the Genomics Research Center at the University of Rochester. None of the authors have any financial conflict of interest related to the research described in this manuscript. This work was supported in part by National Institutes of Health grants R01 HL132899 and R01HL147954 (to P.Y.), start-up funds from Aab Cardiovascular Research Institute of University of Rochester Medical Center (to P.Y.), and American Heart Association Postdoctoral Fellowship 19POST34400013 (to J.W.).

## SI Materials and Methods

### Reagents, antibodies and siRNAs

Human angiotensin II (Cat. No. A9525), E-64d (Cat. No. E8640), 2,3 Butanedione Monoxime (Cat. No. B0753), and 5-Bromo-2’-deoxyuridine (Cat. No. 10280879001) were purchased from Sigma Aldrich, USA. MG132 (Cat. No. 474790) was obtained from Calbiochem, USA. The type II collagenase was purchased from Worthington company (Cat. No. LS004177). Taurin (Cat. No. 1665100) was purchased from Acros organics. In situ cell death detection kit TMR red (Cat. No. 12156792910) was procured from the Roche company. The antifade mounting medium with DAPI (Cat. No. H-1500) was obtained from Vectorlabs company. The DNA plasmids such as Myc- AGTRAP and mCherry-AT1R were purchased from the Sino Biological company. siRNAs including human Fam114a1 (Cat. No. 4392421), and negative control siRNA were procured from ThermoFisher Scientifics and transfected into cardiac fibroblasts by lipofectamine 3000 (ThermoFisher) following the instruction manual.

Antibodies used in this study:

| Primary antibody | Cat No and Company | Dilution | Detection |
| --- | --- | --- | --- |
| Rabbit anti-FAM114A1 | 21638-1-AP; Proteintech | IF (1:200)<br>WB (1:1000) | NIH/3T3; Cardiac fibroblasts |
| Rabbit anti-AGTRAP | 11559-1-AP; Proteintech | WB (1:1000) | NIH/3T3; Cardiac fibroblasts |
| Rabbit anti-AT1R | Ab18801; Abcam | WB (1:1000) | NIH/3T3; Cardiac fibroblasts |
| Rabbit anti-CMTM5 | PA5-62589; ThermoFisher | WB (1:1000) | Heart tissue lysates |
| Rabbit anti-SPG21 | PA5-59176; ThermoFisher | WB (1:1000) | Heart tissue lysates |
| Mouse anti-RAB2B | sc-81921; Santa Cruz | WB (1:500) | Heart tissue lysates |
| Mouse anti-Vimentin | 60330-1; Proteintech | IF (1:300) | Cardiac fibroblasts |
| Mouse anti-GAPDH | 60004-1; Proteintech | WB (1:5000) | Multiple cell types |
| Mouse anti-FLAG | 66008-2; Proteintech | WB (1:1000) | Multiple cell types |
| Mouse anti-α-Tubulin | 66031-1; Proteintech | WB (1:5000) | Multiple cell types |
| Goat anti-Mouse AF488 | A11001; Life technologies | IF (1:500) | Multiple cell types |
| Goat anti-Rabbit AF488 | A11034; Life Technologies | IF (1:500) | Multiple cell types |
| Goat anti-Mouse AF594 | A1105; Invitrogen | IF (1:500) | Multiple cell types |
| Goat anti-Rabbit AF594 | A11012; Invitrogen | IF (1:500) | Multiple cell types |
| Rabbit anti-COL1A1 | SAB2109131; Sigma Aldrich | IF (1:300) | Myofibroblasts |
| Mouse anti-α-SMA | A2547; Sigma Aldrich | IF (1:300)<br>WB (1:1000) | Myofibroblasts |
| Mouse anti-CD45 | BDB56050; BD biosciences | IF (1:200) | Hematopoietic cells |
| Mouse anti-α-Actinin | A7811; Sigma Aldrich | WB (1:5000) | Cardiomyocytes |
| Wheat Germ Agglutinin AF-488 | W11261; Invitrogen | IF (1:1000) | Cardiomyocytes |
| Mouse anti-BrdU | 66241-1; Proteintech | IF (1:200) | Proliferating cells |
| Mouse anti-PDGFRα | AF1062-SP; R&D systems | IF (1:100) | Cardiac fibroblasts |
| Mouse anti-Cardiac Troponin | MA5-12960; ThermoFisher | IF (1:200) | Cardiomyocytes |

### Human tissue samples

All human samples of frozen cardiac tissues, including 20 samples from explanted failing hearts (10 ischemic and 10 dilated cardiomyopathy hearts) and 8 samples from non-failing donor hearts, as well as paraffin section slides from DCM (n=5), ISHF (n=5) and non-failing hearts (n=7) were acquired from the Cleveland Clinic. All surgical procedures and tissue harvesting were performed following the Cleveland Clinic procedures and guidelines. This study was approved by Material Transfer Agreement between the URMC and the Cleveland Clinic. In Figure 1A, total RNA samples from two failing hearts showed degradation during the quality control process and were excluded from RT-qPCR analysis. All human samples were picked up randomly based on the presence or absence of heart failure by our collaborator, Dr. Waihong Wilson Tang at Cleveland Clinic. We are blinded from any clinical data. The identity numbers (IDs) of the frozen cardiac tissues and histology tissues are listed below.

Cardiac Tissue
| Non-Failing ID | Dilated Failing ID | Ischemic Failing ID |
| --- | --- | --- |
| D44 | X2870 | X2854 |
| D156 | X2866 | X2834 |
| D158 | X2860 | X2749 |
| D188 | X2853 | X2679 |
| D204 | X2848 | X2614 |
| D214 | X2846 | X2607 |
| D218 | X2833 | X2590 |
| D247 | X2823 | X2549 |
|  | X2768 | X2445 |

|  | X2755 | X2348 |
| --- | --- | --- |
| **Histology Tissue** |  |  |
| **Non-Failing ID** | **Dilated Failing ID** | **Ischemic Failing ID** |
| D195 | X2670 | X2669 |
| D199 | X2690 | X2732 |
| D214 | X2698 | X2739 |
| D222 | X2675 | X2712 |
| D226 | X2659 | X2679 |
| D237 |  |  |
| D224 |  |  |

### Maintenance of experimental animals and mouse heart failure models

All animal experimental procedures were approved by the University of Rochester Medical Center Animal Care and Use of Committee. In this study, we used WT C57BL/6J and *Fam114a1*^−/−^ (global knockout with the C57BL/6J background) mice generated at Jackson Laboratories (IMPC program). They generated the mice by injecting the Cas9 RNA and four guide sequences ACCAAGCAACCACATCTCCC,AGATGTGGTTGCTTGGTGGT, TTTGACATAGCGTACATTGA, and ATGGAGTTTAACGTTTGCTC, which resulted in a 314 base pair of nucleotides deletion beginning at chromosome 5 positive-strand position 64,995,684 bp, GGGAGATGTGGTTGCTTGGT, and ending after GGAGTTTAACGTTTGCTCAG at 64,995,997 bp (GRCm38/mm10). The mutation deletes exon 3 and 226 bp of flanking intronic sequence including the splice acceptor and donor and is predicted to cause a change of amino acid sequence after residue 116 and early truncation 3 amino acids later. Mice were procured at 8 weeks of age and maintained in vivarium facility and ad libitum free access to standard chow and water.

In this study, we used two mouse heart failure models including angiotensin II (Ang II) osmotic minipump (ALZA Corporation, CA) implantation and myocardial infarction (MI) surgery (left anterior descending coronary artery ligation). In the entire study, we used age-matched male mice at the age of ∼8-12 weeks. All the mouse surgeries were done by the mouse Microsurgical Core facility at URMC. We used 6-8 mice from each genetic background for individual treatment group to get the statistical significance of different groups of the study.

For the osmotic minipump implantation, WT and *Fam114a1*^+/–^ male mice from sibling mating at 8–12 weeks of age were subjected to subcutaneous infusion with vehicle saline or Ang II for 4 weeks using osmotic mini-pumps in a randomized and blinded manner by microsurgeons from the microsurgical core facility of Aab CVRI. Mice were anesthetized using 2.0% isoflurane and placed on a heated surgical board. A side/upper back area skin incision was made, and the mini-osmotic pump was inserted subcutaneously and set to deliver Ang II or vehicle at a rate of 1.4 mg/Kg/day. The incision was then closed with 6-0 coated vicryl in a subcuticular manner, and the animals were allowed to recover. The sutures were removed 2 weeks after the pumps were transplanted. The pumps were not removed and remained for a period of 4 weeks. The animals were euthanized after 4 weeks of Ang II infusion and mouse hearts were harvested for experiments, including RNA, protein extraction and sectioning.

The LAD ligation-based MI surgery was performed as previously described by the Mouse Microsurgical Core of Aab CVRI^1^. For MI surgery, male or female mice were placed on a heating pad and the airway stabilized by endotracheal intubation and mechanical ventilation provided (inspiratory tidal volume of 250 μL at 130 breaths/min). The mice were given SR Buprenorphine 2.5 mg/Kg via subcutaneous injection. Isoflurane flow was continually maintained approximately at 1.5% along with oxygen. A midline cervical incision was made to expose the trachea for intubation with a PE90 plastic catheter. The catheter was connected to a Harvard mini vent supplying supplemental oxygen with a tide volume of 225-250 μl and a respiratory rate of 130 strokes/min. Surgical plane anesthesia was subsequently maintained at 1-1.5% isoflurane. The skin was incised and the chest cavity opened at the level of the 4th intercostal space. Oral intubation was employed by placing PE 90 tubing in the mouth and advancing slowly into the trachea. Mechanical P.I. ventilation (tidal volume of approximately 0.4 ml at 130 breaths/min) was then begun. After intubation, a midline incision was made between the sternum and the left internal mammalian artery. Alternatively, a lateral incision (left thoracotomy) was made in the fourth intercostal space. The mouse heart was exposed and the left coronary artery branch points are visualized under 10x magnification before ligation, and the LAD coronary artery ligated intramurally 2 mm from its ostial origin for standard MI with a 9-0 proline suture. Transmural ischemia was assured by color loss on the left ventricle wall and ST-segment elevation noted on the electrocardiogram. Lungs were inflated and the chest was closed in two layers; the ribs (inner layer) were closed with 6-0 coated vinyl sutures in an interrupted pattern. The skin was closed using 6-0 nylon or silk sutures in a subcuticular manner. The anesthesia was stopped and once the mouse was breathing on its own, the mouse was removed from the ventilator and allowed to recover in a clean cage on a heated pad. A sham operation was performed using the same procedure, but a suture was passed under the LAD coronary artery without ligation.

For mouse experiments, age/sex/genetic background matched mice were randomly separated into indicated groups. Animal operations, including Ang II infusion, MI surgery, and echocardiography measurement, were performed blindly by the Microsurgical Core surgeons. Heart sections were prepared by the Histology Core. For group size justification, we have performed a power analysis both G*power version 3.1.9.6. The assumptions include the same standard variance in each study group, effect 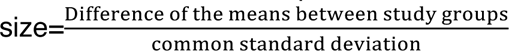, alpha level=0.05, power=0.9 and number of study groups. The effect size for specific experiments is assumed based on previous similar studies or literature. As an exemplary experiment, the standard deviation for weight after MI or Ang II treatment is about 10%. The minimum difference to be considered as significance is 25% in MI- or Ang II-induced cardiac hypertrophy and heart weight gain. With an overall type I error rate (alpha level) of 5%, at least 5 mice per treatment group are required to achieve 90% power to detect the difference of heart weight. In previous experiences from our Microsurgical Core, we have observed a survival rate of ∼90% after the MI procedure. To offset the possible loss of one mouse per treatment, we used at least 6 mice per treatment group. In rare cases, mice might die after surgery which might reduce the number of mice.

### Echocardiography

For Ang II infusion mouse model, M-mode short axis echocardiographic image collection was performed using a Vevo2100 echocardiography machine (VisualSonics, Toronto, Canada) and a linear-array 40 MHz transducer (MS-550D). Heart rate was monitored during echocardiography measurement. Image capture was performed in mice under general isoflurane anesthesia with heart rate maintained at 500-550 beats/min. LV systolic and diastolic measurements were captured in M-mode from the parasternal short axis. Fraction shortening (FS) was assessed as follows: % FS = (end diastolic diameter - end systolic diameter) / (end diastolic diameter) x 100%. Left ventricular ejection fraction (EF) was measured and averaged in both the parasternal short axis (M-Mode) using the tracing of the end diastolic dimension (EDD) and end systolic dimension (ESD) in the parasternal long axis: % EF=(EDD-ESD)/EDD. Hearts were harvested at multiple endpoints depending on the study. In addition to EF and FS, left ventricular end-diastolic diameter (LVEDD), left ventricular end-systolic diameter (LVESD), wall thickness of left ventricular anterior (LVAWT) and posterior (LVPWT) were also assessed. For MI mouse model, B-mode long axis echocardiographic imaging was performed and left ventricle EF, FS, end systolic volume (LVESV), and end diastolic volume (LVEDV) were measured.

### Adult cardiomyocyte and cardiac fibroblasts isolation and culturing

Langendorff perfusion system was used to isolate adult cardiomyocytes (CMs) and cardiac fibroblasts from the murine heart. Mice were fully anesthetized via intraperitoneal injection of ketamine/xylazine. Once losing pedal reflex, the mouse was secured in a supine position. The heart was excised and blood was removed using perfusion buffer. The heart was then fastened onto the CM perfusion apparatus where perfusion was initiated using the Langendorff mode. Our Langendorff perfusion and digestion consisted of three steps at 37°C: 4 mins with perfusion buffer (0.6 mM KH2PO4, 0.6 mM Na2HPO4, 10 mM HEPES, 14.7 mM KCl, 1.2 mM MgSO4, 120.3 mM NaCl, 4.6 mM NaHCO3, 30 mM taurine, 5.5 mM glucose, and 10 mM 2,3-butanedione monoxime), then switched to digestion buffer (300 U/ml collagenase II [Worthington] in perfusion buffer) for 3 mins, and finally perfused with digestion buffer supplemented with 40 μM CaCl2 for 8 mins. After perfusion, the ventricle was placed in sterile 35 mm dish with 2.5 ml digestion buffer and shredded into several pieces with forceps. 5 ml stopping buffer (10% FBS, 12.5 μM CaCl2 in perfusion buffer) was added and pipetted several times until tissues disperse readily, and solution turned cloudy. The cell solution was passed through 100 μm strainer. CMs were settled by incubating the cell suspension at 37°C for 30 mins. The CMs were resuspended in 10 ml stopping buffer and subjected to several steps of calcium ramping: 100 μM CaCl2, 2 mins; 500 μM CaCl2, 4 mins; 1.4 mM CaCl2, 7 mins. Then the CMs were seeded onto a glass bottom dish (Nest Biotechnology) pre-coated with laminin (ThermoFisher). Plates were centrifuged for 5 mins at 1,000 g at 4°C to increase the adherence, cultured at 37°C for ∼1 hr, and then switched to CM media (MEM [Corning] with 0.2% BSA, 10 mM HEPES, 4 mM NaHCO3, 10 mM creatine monohydrate, 1% penicillin/streptomycin, 0.5% insulin-selenium-transferrin and blebbistatin) for cell culture and further treatments.

Cardiac fibroblasts (CFs) from the supernatant were pelleted for 5 mins at 1,200 rpm at 4°C. CFs were plated in 4-5 ml CF media (DMEM with 10% FBS and 1% penicillin/streptomycin) in 60 mm plate and washed vigorously 3-5 times with 2 ml 1x PBS for several times after 2-3 hrs to remove the unattached cells and debris, and replaced with fresh CF media. For CF only isolation, pre- weened mice were fully anesthetized and the heart was directly cut into small pieces and digested in the digestion buffer for 4x 10 mins at 37°C with slow stirring and CFs were plated the same as Langendorff isolation of CFs.

### RNA isolation and RT-qPCR

For heart tissues (human and mouse) or cell samples, the RNA extraction was performed using TRIzol reagent (ThermoFisher) following instructions in the manual and used for the detection of the expression of specific genes. Briefly, the tissues were homogenized in TRIzol using Minilys Personal Homogenizer (Bertin Technologies) and placed on ice for 15 mins to lyse the tissue. Genomic DNA was removed using DNase I treatment followed by the phenol-chloroform-isoamyl alcohol extraction method. For the mRNA detection, 1 μg of total RNA was used as a template for reverse transcription using the iScript cDNA Synthesis Kit (Bio-Rad). RT-qPCR was performed with cDNA, primers of specific targets of interest, and IQ SYBR Green Supermix (Bio-Rad). Data were analyzed using the formula of the ΔΔC(t) method. cDNA was used for detecting the expression of Fam114a1, Agtrap, At1r, and the marker genes, Nppa, Nppb, Col1a1 and Col3a1. 18S rRNA or Gapdh was used as a normalization control for mRNA expression. The SYBR Green primer sequences or the Taqman probes were listed as below.

SYBR green qPCR procedure: 1) initial denaturation at 95°C for 60 seconds. 2) 40 cycles of denaturation at 95°C for 10 seconds and annealing/extension at 60°C for 45 seconds. 3) melt curve analysis by 0.5°C increments at 5 seconds/step between 65-95°C. Taqman assay qPCR procedure: 1) initial denaturation at 95°C for 10 mins. 2) 40 cycles of denaturation at 95°C for 15 seconds and annealing/extension at 60°C for 60 seconds.

qPCR primers used in this study (SYBR green and Taqman probes):

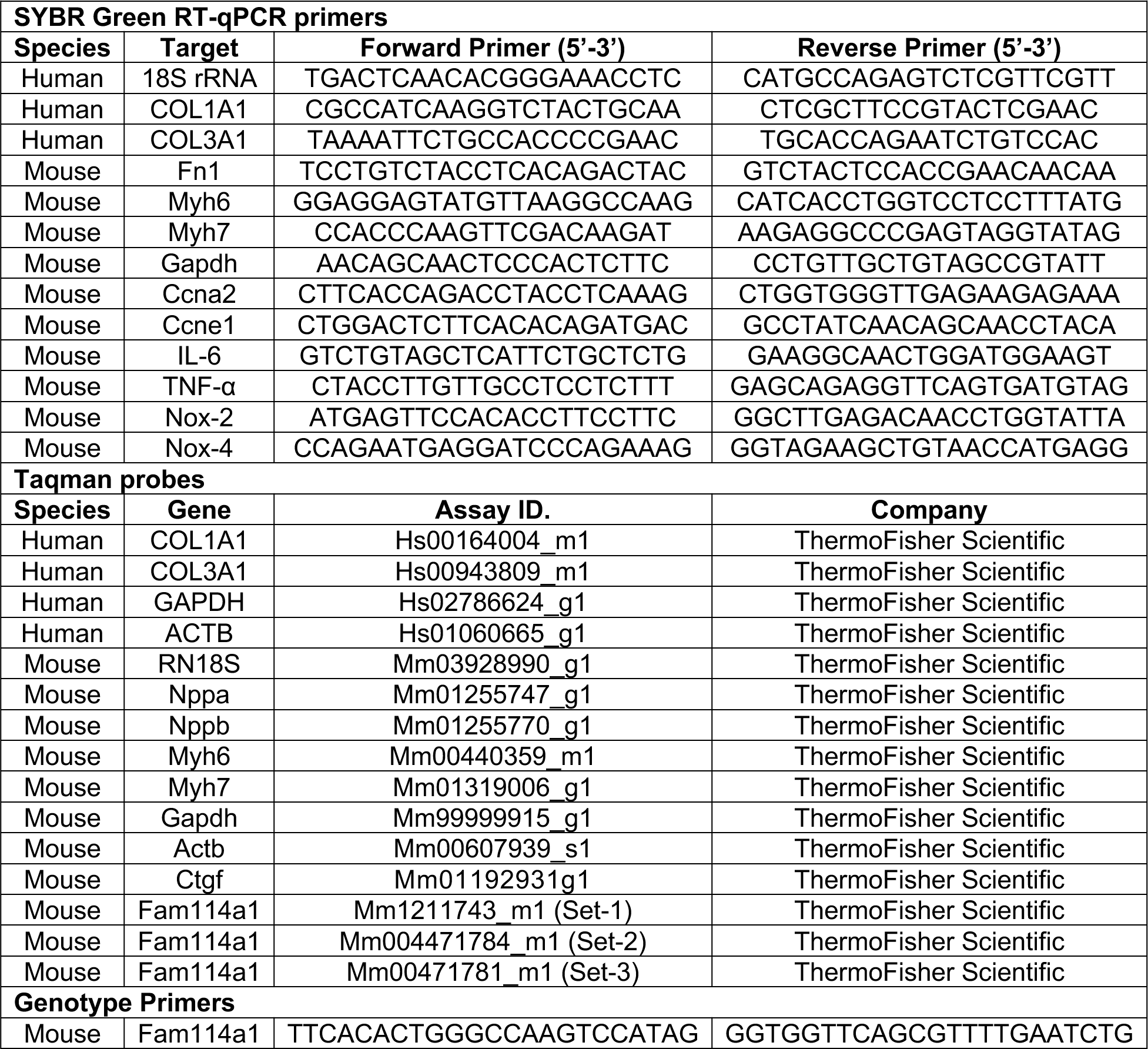
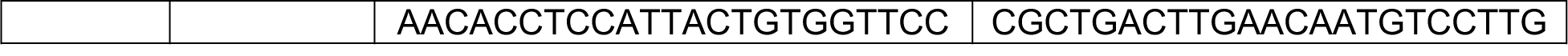

### Immunofluorescence staining of Heart tissue sections and CFs

Mice were sacrificed and hearts were immediately removed, washed in ice-cold PBS, and fixed in 10% neutral buffered formalin, and processed for paraffin embedded sections in the Histological Core of Aab CVRI. Tissue sections were cut at a cross-section of 5 μm thickness 250 μm intervals were used for immunohistochemical analysis and to quantify the scar area. For immunofluorescence, paraffin-embedded slides were deparaffinized in a series of xylenes as followed by 3 mins of incubations in 100% ethanol, 95% ethanol and then placed in a distilled water. Antigen retrieval was performed in citric acid buffer (pH 6.0) followed by quenching in 3% H2O2 in PBS for 30 mins at RT. Sections were blocked in blocking buffer (2% BSA, 0.5% Triton X-100, 5% goat serum) for 2 hrs. Then slides were incubated with primary antibodies (as listed in the antibody table) for overnight at 4°C. After primary antibody incubation, slides were washed in 1x PBS followed by the secondary antibody (AlexFluor-488 or AlexFluor-594 conjugated) incubation in blocking solution for 1 hr at room temperature (RT). Slides were then washed with 1x PBS (3x 5 min) and mounted with DAPI (Vectorlabs), covered by coverslips and air-dried (or kept in PBS buffer inside before imaging), and the images obtained using the Olympus FV1000 confocal microscope and the intensity was measured by NIH Image J software. Four sections for each condition were used and for each section 5-7 randomized fields of images were captured (4 sections x 7).

### Wheat germ agglutinin (WGA) staining

WGA staining was used to quantify the size of CMs in the murine heart. Deparaffinization, antigen retrieval and quenching of auto-fluorescence were performed as described above. Heart tissue sections of WT and *FAM114A1*^−/−^ mice from different treatments were probed with 10 μg/ml WGA-Alexa Fluor-488 (ThermoFisher) to stain the cardiomyocyte membrane for 1 hr at RT and followed by 3x 5 mins washes with 1x PBS. The slides were covered by coverslips with antifade solution (containing DAPI) for imaging. Cardiomyocytes were measured in the whole heart of vehicle and Ang II infused mice and in remote and border zone areas of hearts of MI mice. The images were taken in the fluorescence microscope and cross-sectional areas were quantified and measured using Image J Version 2.0. software (NIH, USA) using the hand drawing tool to outline the myocytes. Myocyte size was taken from images of at least 3-4 fields per heart and in total 300- 400 cells were measured.

### Terminal deoxynucleotidyl transferase dUTP nick end labeling (TUNEL) staining

The tissue sections were washed with PBS twice and fixed using 4% paraformaldehyde for 20 mins. Cells were permeabilized with 0.5% of Triton X-100 for 5 mins, and incubated in TUNEL reaction mixture (In Situ Cell Death Detection Kit; Sigma, 11684795910) for 1 hr at 37°C in the dark. Finally, cells were washed with PBS for 3x 5 mins, air dried, and mounted with DAPI-containing antifade medium. Images were captured using a BX51 microscope (Olympus).

### Cardiac fibroblasts migration assay

To determine the cardiac fibroblasts migration ability, 5x 10^4^ primary isolated ACFs from WT and FAM114A1^−/−^ were plated per well 24 well plates. Fibroblasts were stimulated with human angiotensin II (100 nM) or unstimulated (vehicle). Fibroblast monolayers were then scratched with a 200 μl pipet tip. To analyze the migration of the fibroblasts, the same scratched area was captured with the Olympus microscope after 0, 6,12, and 24 hrs. The migration ration ratio was calculated as (cell-free area at 0 hr – cell-free area at 6,12 or 24 hr) / cell-free area at 0 hr. The cell-free area was calculated using the Image J software (NIH, USA). The migration assay was carried out in 3 biological replicates with 3 wells per each treatment condition.

### Triphenyl Tetrazolium Chloride (TTC) staining for measurement of infarct size

Excised hearts were perfused with 1x PBS to remove the blood, and sectioned into 4-5 levels (2 mm thick). The sliced hearts were placed in a petri dish with 1% TTC in 1x PBS and incubated for 15 mins at 37°C. Then, tissue slices were fixed with 10% formalin for 1 hr. Hearts were visualized using a bright field microscope. Quantification of infarct size was performed using Image J Version 2.0. software (NIH, USA) by normalizing total scar (white color area) to the left ventricle wall (% LV free wall) and averaging across four cross-sectional levels of the heart (apex to ligature).

### Mouse Cytokine Array

Blood was collected from the mice through the cardiac puncture and placed in collection tubes coated with EDTA (Mini Collect, Greiner Bio-One). The samples were centrifuged at 1,200 rpm for 10 mins, and serum was collected from each sample and stored at -80°C until ready to use. A total of 100 μl of serum was used by combining serum from 5 animals from each treatment of WT and *FAM114A1*^−/−^ mice. Proteome profiler mouse cytokine array (ARY006, R&D Systems) assay was performed according to the manufacturer’s instructions. Cytokines were detected by chemiluminescence with exposure up to 1-5 mins using the ChemiDoc Imaging System (Bio-Rad). The relative intensity of the signals was calculated using Image Lab software (Bio-Rad).

### Immunoprecipitation and Western blot analysis

Heart tissues, isolated CMs, and cultured CFs were homogenized in ice-cold RIPA lysis buffer along with protease inhibitor cocktails (Santa Cruz). Cell debris was removed by centrifugation for 10 mins at 10,000 rpm, 4°C. Total protein concentration was determined by Bradford assay (Bio-Rad). An equal amount of protein was loaded on to 10% and 12% SDS-PAGE gels and then transferred to PVDF membranes. The membranes were blocked in the 5% milk in PBST buffer for 1 hr at RT. The respective membranes were probed with specific primary antibodies for target proteins in 4% BSA (Sigma Aldrich) in PBST buffer for overnight. After several washes with the PBST buffer, the blots were incubated with a horseradish peroxidase-conjugated secondary antibody in 3% milk with PBST buffer for 1 hr and developed using the ECL reagent (Bio-Rad).

### RNA-Seq data processing and alignment

Total RNA extracted from CF cells isolated from WT and *Fam114a1*^−/−^ male mice were treated with DNase I (NEB) to remove potential genomic DNA in the RNA samples. The DNase I treated RNA samples were purified with phenol:chloroform:isoamyl alcohol and then subjected to RNA-Seq at the Genomic Research Center of URMC. Raw reads generated from the Illumina HiSeq2500 sequencer were demultiplexed using bcl2fastq version 2.19.0. Quality filtering and adapter removal were performed using Trimmomatic version 0.36^2^ with the following parameters: “TRAILING:13 LEADING:13 ILLUMINACLIP: adapters.fasta:2:30:10 SLIDINGWINDOW:4:20 MINLEN:15”. Processed/cleaned reads were then mapped to the *Mus musculus* reference genome (GRCm38, mg38) with STAR_2.5.2b^3^ given the following parameters: “—twopassMode Basic --runMode alignReads --genomeDir ${GENOME} --readFilesIn ${SAMPLE} --outSAMtype BAM SortedByCoordinate --outSAMstrandField intronMotif --outFilterIntronMotifs RemoveNoncanonical”. The subread-1.5.0^4^ package (featureCounts) was used to derive gene counts given the following parameters: “-s 2 -t exon -g gene_name” and the gencode M12 gene annotations. Differential expression analysis and data normalization was performed using DESeq2-1.16.1^5^ with an adjusted p-value threshold of 0.05 within an R-3.4.1^6^ environment. A batch factor was given to the differential expression model in order to control for batch differences. Gene ontology and KEGG pathway enrichment analyses were performed using DAVID Bioinformatics Resources 6.8^7^.

## Supplemental figures

**Figure S1.**
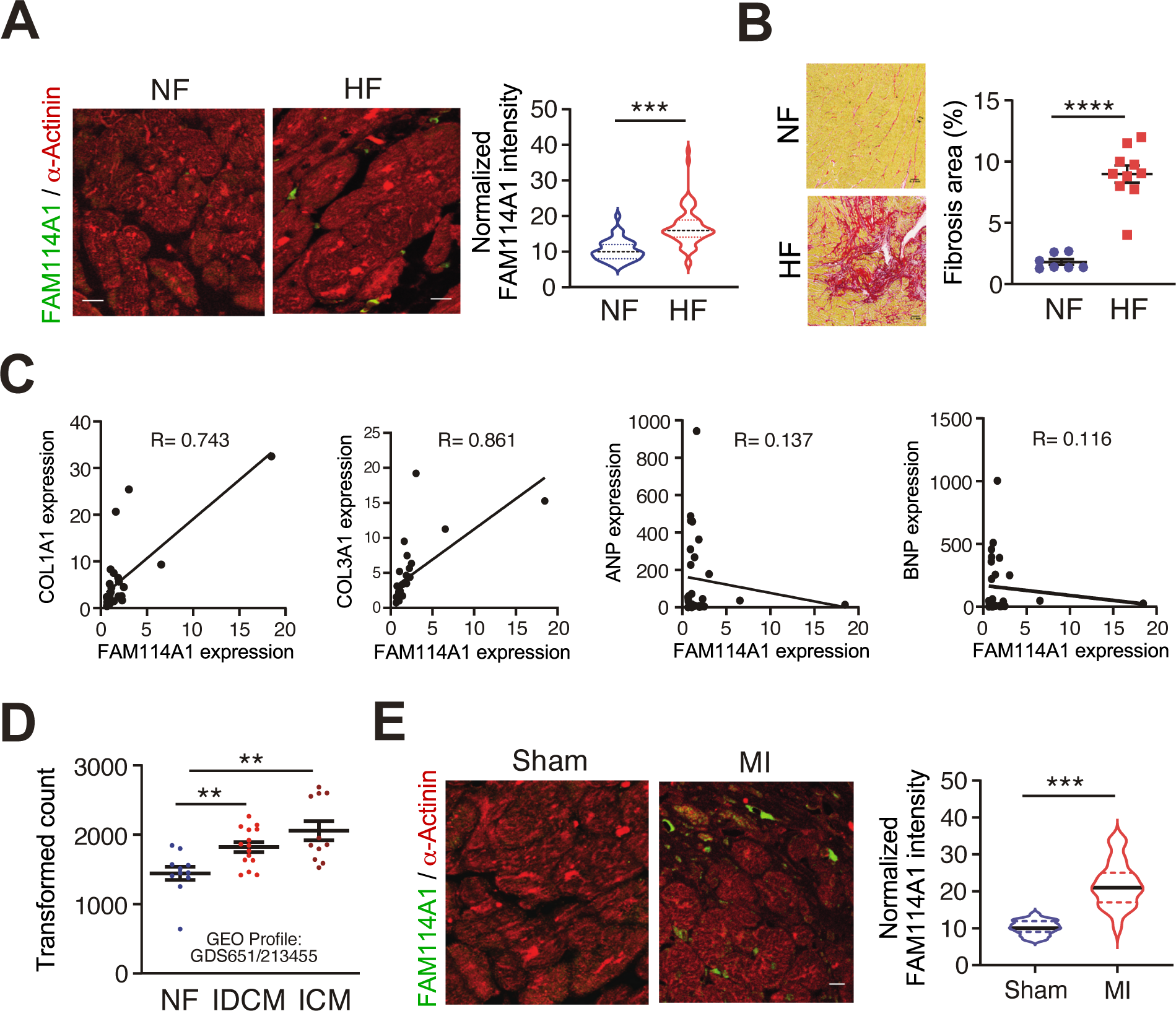
Increased FAM114A1 expression in human and mouse heart failure and correlation with collagen expression. (A) IF analysis and quantification of FAM114A1 protein expression in human failing hearts. *α*- actinin IF indicates FAM114A1 protein expression in non-myocyte cells such as CFs (100-130 cells were quantified). (B) Picrosirius red staining of human failing hearts versus non-failure donor hearts (n=7 for NF; n=10 for HF). (C) FAM114A1 expression is correlated with the expression of collagens (not ANP or BNP) in human heart samples (n=26; 8 for NF and 18 for HF). Pearson correlation coefficient was presented. 18S rRNA was used as a normalizer. (D) *FAM114A1* mRNA expression is increased in idiopathic dilated cardiomyopathy (IDCM, n=15) and ischemic cardiomyopathy (ICM, n=11) patients compared to non-failure (NF, n=11) donor heart tissues from a publicly available microarray dataset GDS651 / 213455_at. (E) IF analysis and quantification of FAM114A1 protein expression in mouse MI-derived failing hearts. *α*-actinin IF indicates FAM114A1 protein expression in non-myocyte cells such as CFs (n=4 per group).100-150 cells were counted for the quantification, Data were represented as mean±SEM. Statistical significance was confirmed by unpaired two-tailed Mann Whitney test for data not normally distributed for A, unpaired two-tailed Student *t* test for D, E.

**Figure S2.**
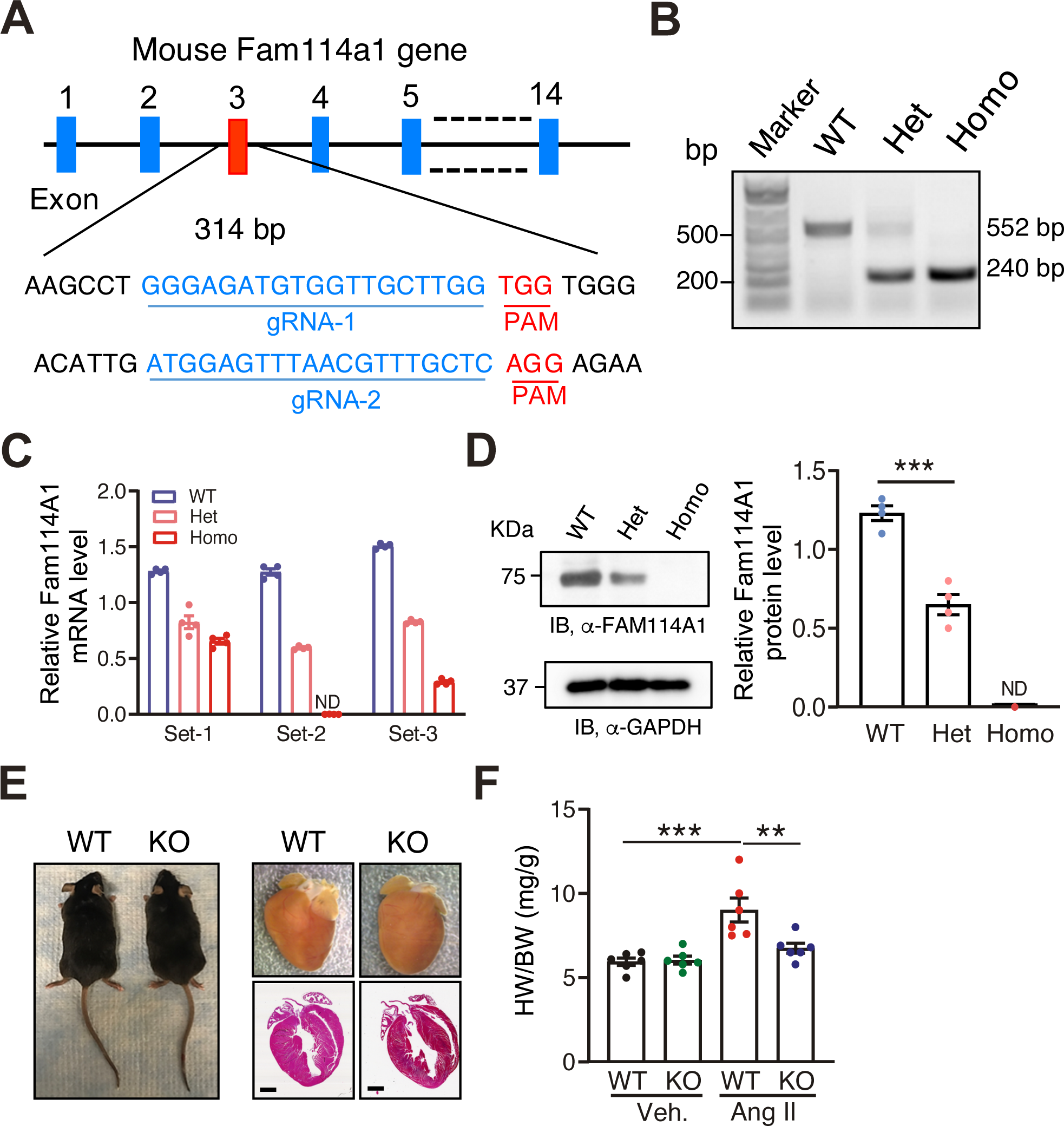
Characterization of *Fam114a1*^−/−^ mice at baseline. (A) Schematic of the generation of *Fam114a1*^−/−^ mice by CRISPR technology from IMPC. (B) Genotyping of WT (wild type), *Fam114a1*^+/–^ (het: heterozygote)*, and Fam114a1*^−/−^(homo: homozygote) mice. (C) RT-qPCR detection of *Fam114a1* mRNA using different primers spanning exon 1 and 2 (Set-1), exon 2-3 (Set-2), and exon 5-6 (Set-3). N=4 per each group. (D) Immunoblot shows FAM114A1 protein expression in the hearts of WT, *Fam114a1*^+/–^*, and Fam114a1*^−/−^ mice. Quantification of FAM114A1 protein intensity is shown. N=4 per each group. (E) Body size and heart morphology of WT and *Fam114a1*^−/−^ mice at baseline. (F) *Fam114a1*^−/−^ mice show reduced HW/BW (heart weight/body weight) ratio after 4 weeks of Ang II infusion compared to WT mice. n=6 for each group. Data were represented as mean±SEM. Statistical significance was confirmed by unpaired two-tailed Student *t* test for D and two-way ANOVA with Tukey’s multiple comparisons test for F.

**Figure S3.**
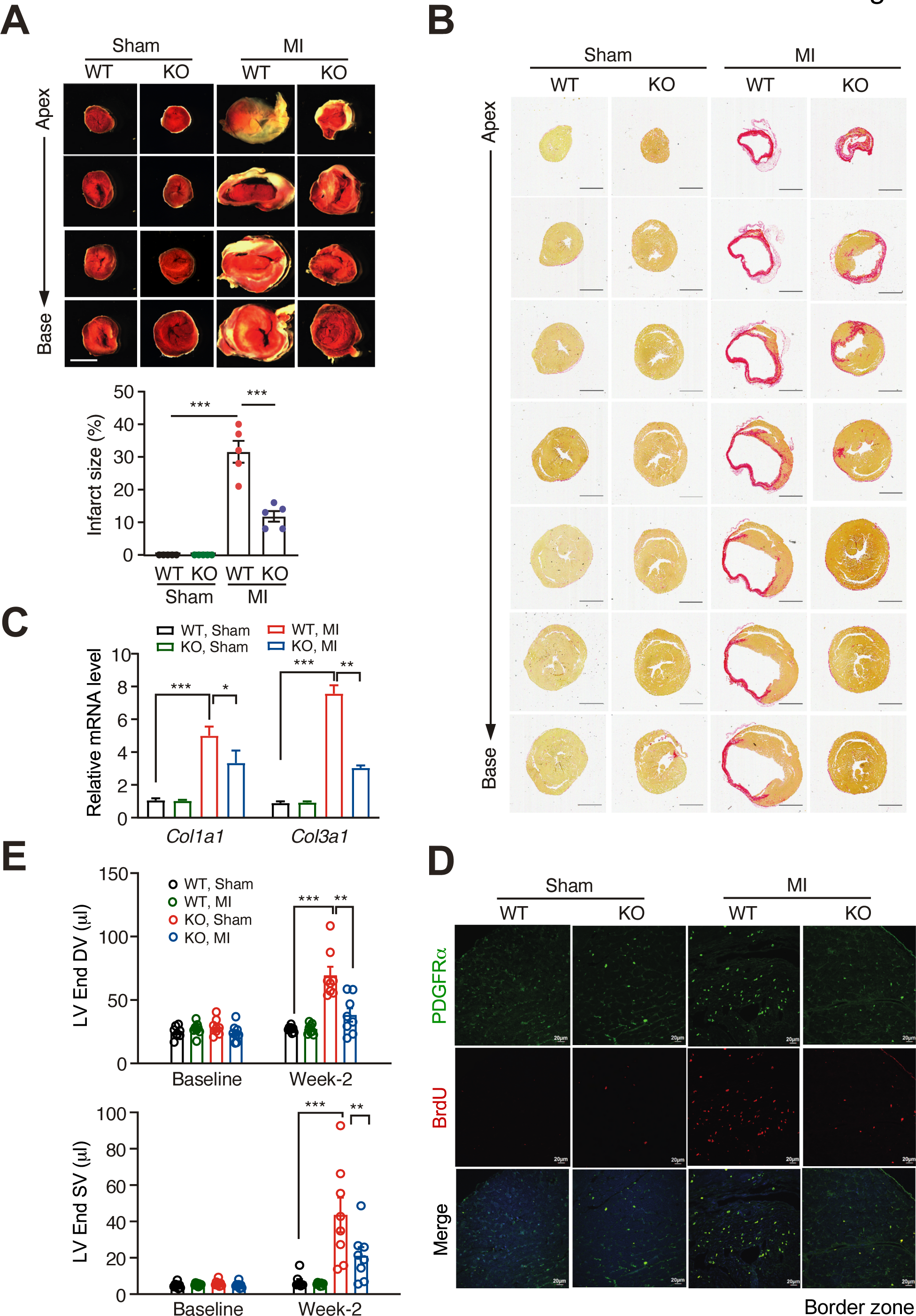
Phenotypic characterization of *Fam114a1*^−/−^ mice after MI. (A) TTC (2,3,5-Triphenyltetrazolium chloride) staining of hearts of WT and *Fam114a1*^−/−^ mice after 20 days post Sham or MI surgery (n=5 per group). The infarct area is calculated from apex to base of the heart sections. (B) Representative images of Picrosirius red staining of cross sections of heart tissues of WT and *Fam114a1^−/−^* mice after 20 days post MI surgery. N=5 for each group. Scale bar: 1 mm. (C) RT-qPCR measurement of cardiac fibrosis marker gene expression in WT and *Fam114a1^−/−^* hearts after MI or Sham surgery. 18S rRNA was used as a normalizer. N=3 for each group. (D) Cardiac fibroblast cell proliferation was measured using BrdU/PDGFR-α staining in the mouse heart tissue sections after 20 days post MI or Sham surgery (n=4). 100-150 cells were counted for the quantification in the border zone. (E) Echocardiography measurement of end diastolic and systolic volume in WT and *Fam114a1*^−/−^ mice after 2 weeks post sham or MI surgery. N=8 for each group. Data were represented as mean±SEM. Statistical significance was confirmed by two-way ANOVA with Tukey’s multiple comparisons test for A, C, E.

**Figure S4.**
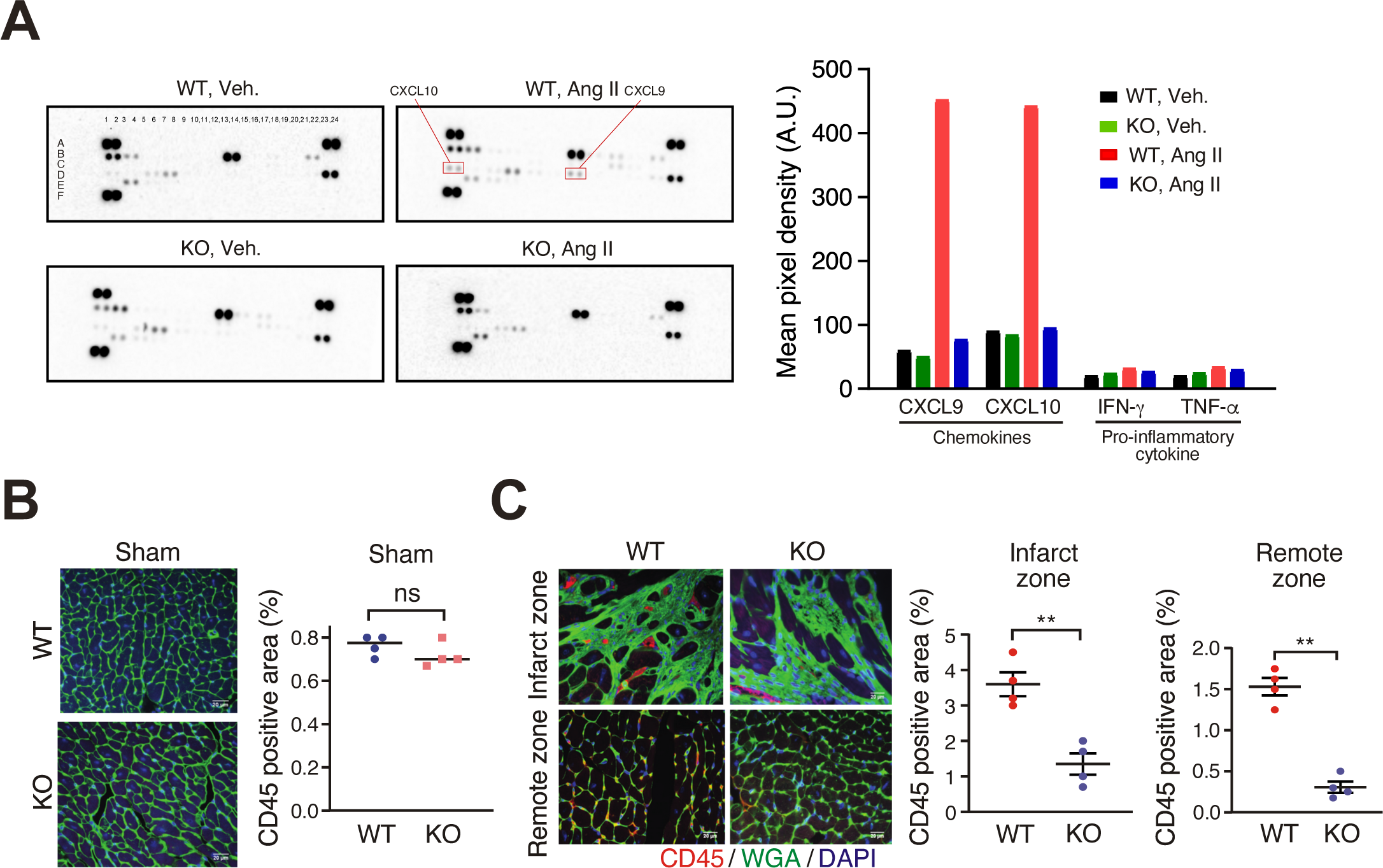
Cytokine production and immune cell infiltration in *FAM114A1*^−/−^ mice after MI. (A) Cytokine array analyses of serum samples of WT and *Fam114a1^−/−^* mice after vehicle or Ang II infusion. N=3 for each group. Three samples were pooled together for analysis. (B) CD45 IF (red) and WGA (green) staining in WT and *Fam114a1*^−/−^ mice after 20 days post Sham surgery. N=4 per group. (C) CD45 IF (red) and WGA (green) staining in the remote and infarct area of WT and *Fam114a1*^−/−^ hearts after 20 days post MI surgery. N=4 per group. Data were represented as mean±SEM. Statistical significance was confirmed by unpaired two-tailed Student *t* test for B, C.

**Figure S5.**
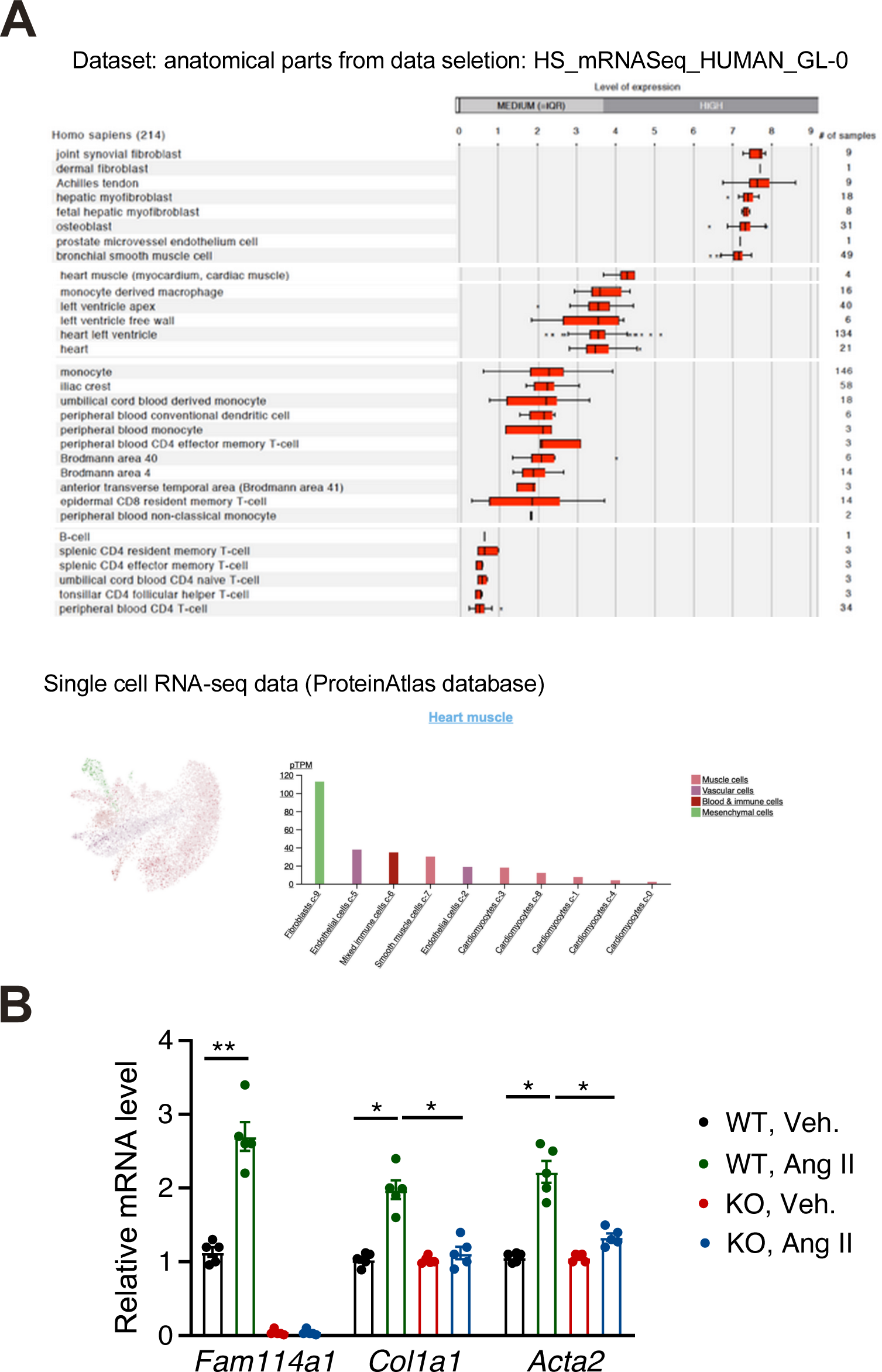
FAM114A1 deletion reduces myofibroblast marker gene expression upon Ang II stimulation. (A) Upper panel: The Genevestigator database provides expression ranking of FAM114A1 across multiple human cell types. Lower panel: The Protein Atlas database provides expression ranking of *FAM114A1* mRNA across different cardiac cell types in the human heart. (B) RT-qPCR measurement of *Fam114a1* and myofibroblast activation marker genes in Ang II or vehicle treated primary CFs isolated from WT and *Fam114a1*^−/−^ mice. 18S rRNA was used as a normalizer (n=5). Data were represented as mean±SEM. Statistical significance was confirmed by two-way ANOVA with Tukey’s multiple comparisons test for B.

**Figure S6.**
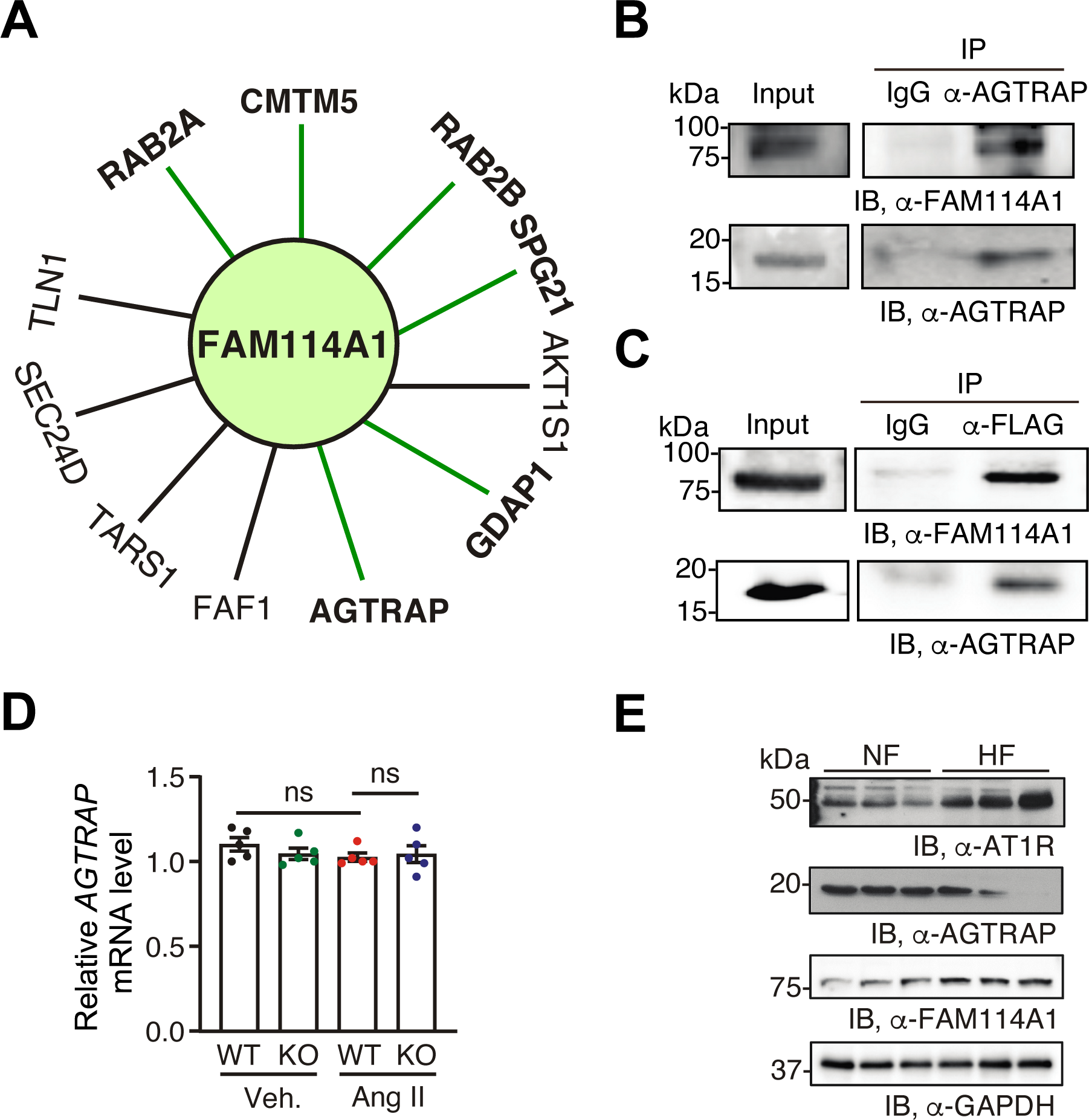
FAM114A1 interacts with AGTRAP. (A) Candidate interacting proteins of FAM114A1 from publicly available databases (BioGRID and BioPlex). Proteins in bold were discovered by yeast two hybrid screen while others were found by co-fractionation and mass spectrometry. (B) Immunoprecipitation and immunoblot confirm the interaction of FAM114A1 with AGTRAP using AGTRAP antibody in isolated primary mouse CFs from WT mice. (C) Immunoprecipitation and immunoblot confirm the interaction of FLAG-FAM114A1 with AGTRAP using FLAG antibody in mouse NIH/3T3 fibroblast cells. (D) *Agtrap* mRNA expression in vehicle or Ang II treated (24 hrs) primary mouse CFs from WT and *Fam114a1*^−/−^ mice. (E) Protein expression of endogenous FAM114A1, AGTRAP, and AT1R in human failing hearts compared to non-failure donor hearts by Western blot analysis. Data were represented as mean±SEM. Statistical significance was confirmed by two-way ANOVA with Tukey’s multiple comparisons test for D.

**Figure S7.**
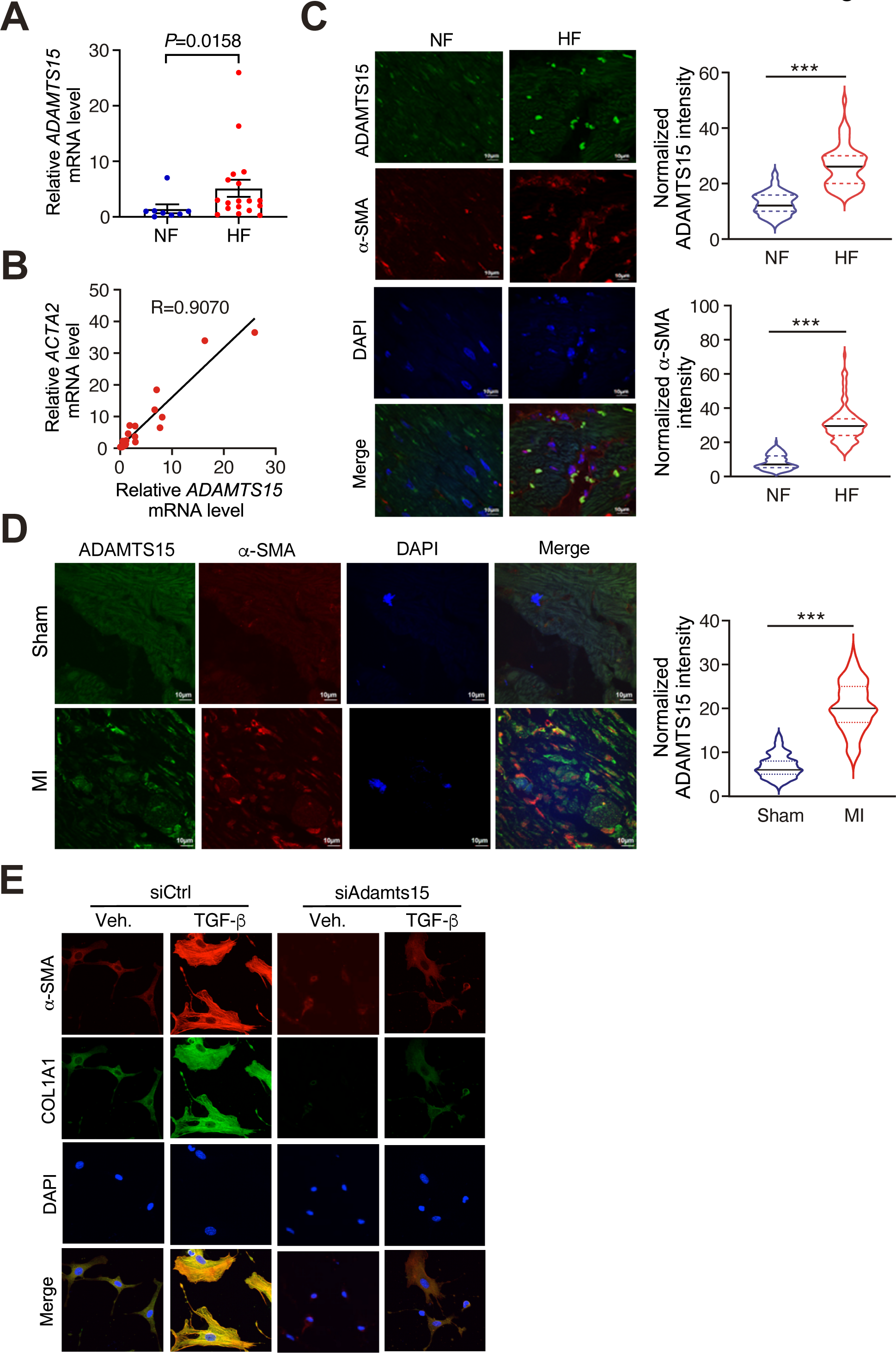
ADAMTS15 upregulation is associated with human and mouse cardiac fibrosis and heart failure. (A) *ADAMTS15* mRNA expression in failing human hearts (n=18) compared to non-failure donor hearts (n=8). 18S rRNA was used as a normalizer. (B) *ADAMTS15* expression is correlated with the expression of *ACTA2* in human heart samples (n=26; 8 for NF and 18 for HF). Pearson correlation coefficient was presented. 18S rRNA was used as a normalizer. (C) FAM114A1 protein expression is increased in failing human hearts (n=5) compared to non-failure hearts (n=5). ∼100-130 cells with positively signals located in MI infarct areas were counted for the quantification. (D) IF analysis and quantification of FAM114A1 protein expression in MI (20 days post-surgery) treated mouse heart tissue sections after Sham or MI surgery (n=6 for both groups). 100-150 cells per heart were counted for the quantification. Scale bar: 10 μm. (E) Representative images of COL1A1 and *α*-SMA IF in TGF-*β*-treated primary mouse CFs after knockdown of *Adamts15*. N=100-120 cells from three biological replicates were analyzed. Representative images were shown. Data were presented as mean±SEM. Comparisons of means between two groups were performed by unpaired two-tailed Mann Whitney test for data not normally distributed for A and unpaired Student *t* test for data normally distributed for C, D.

**Figure S8.**
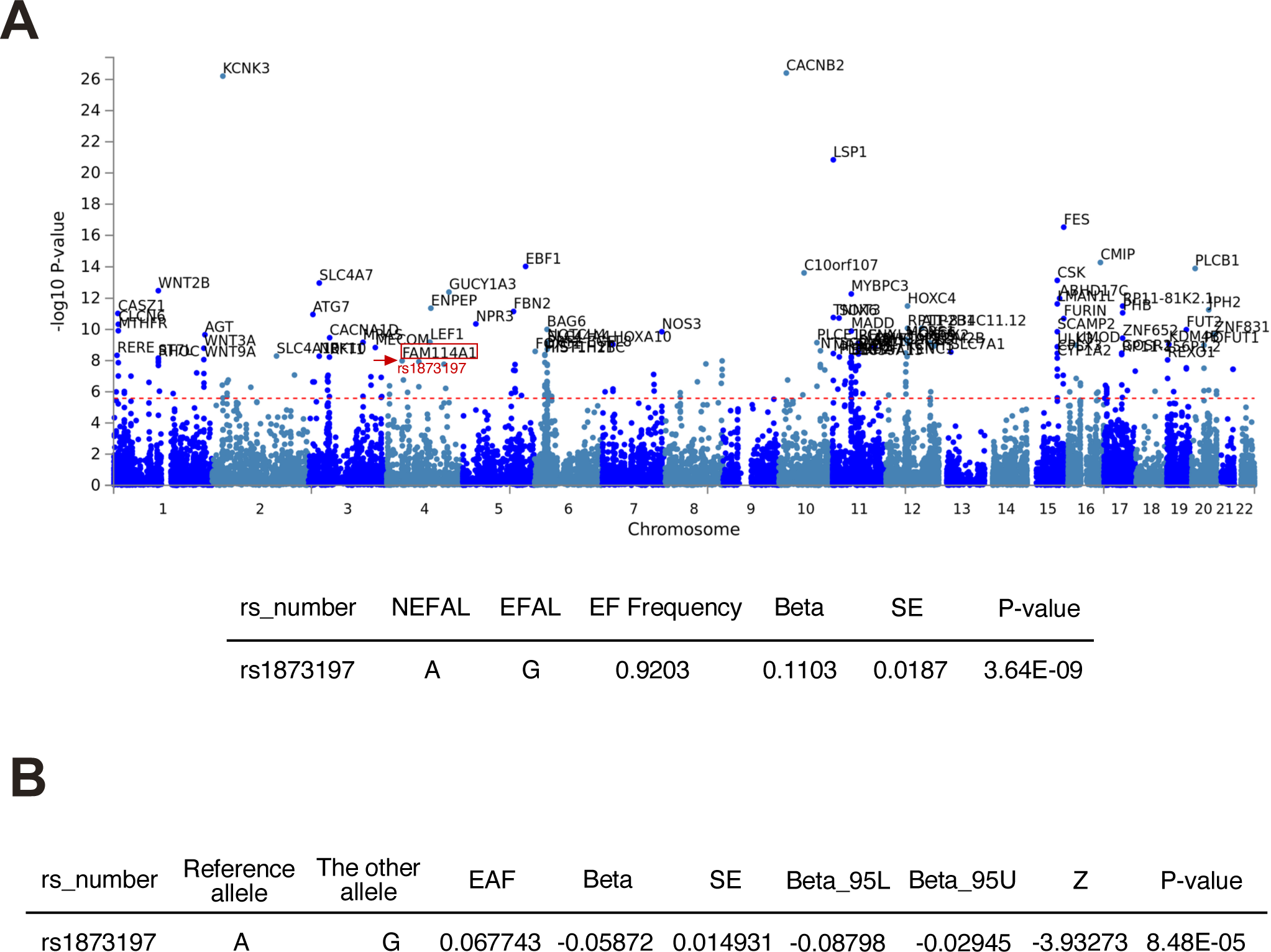
Genome wide association of FAM114A1 SNP rs1873197 with human coronary artery disease and myocardial infarction. (A) Manhattan plot of human genome-wide association study containing FAM114A1 SNP rs1873197 associated with CAD (from GWASATLAS database). (B) FAM114A1 SNP rs1873197 as a human genomic locus associated with myocardial infarction (n=639,000 human MI subjects and n=192,000 human coronary atherosclerosis subjects).

**Table S1.**
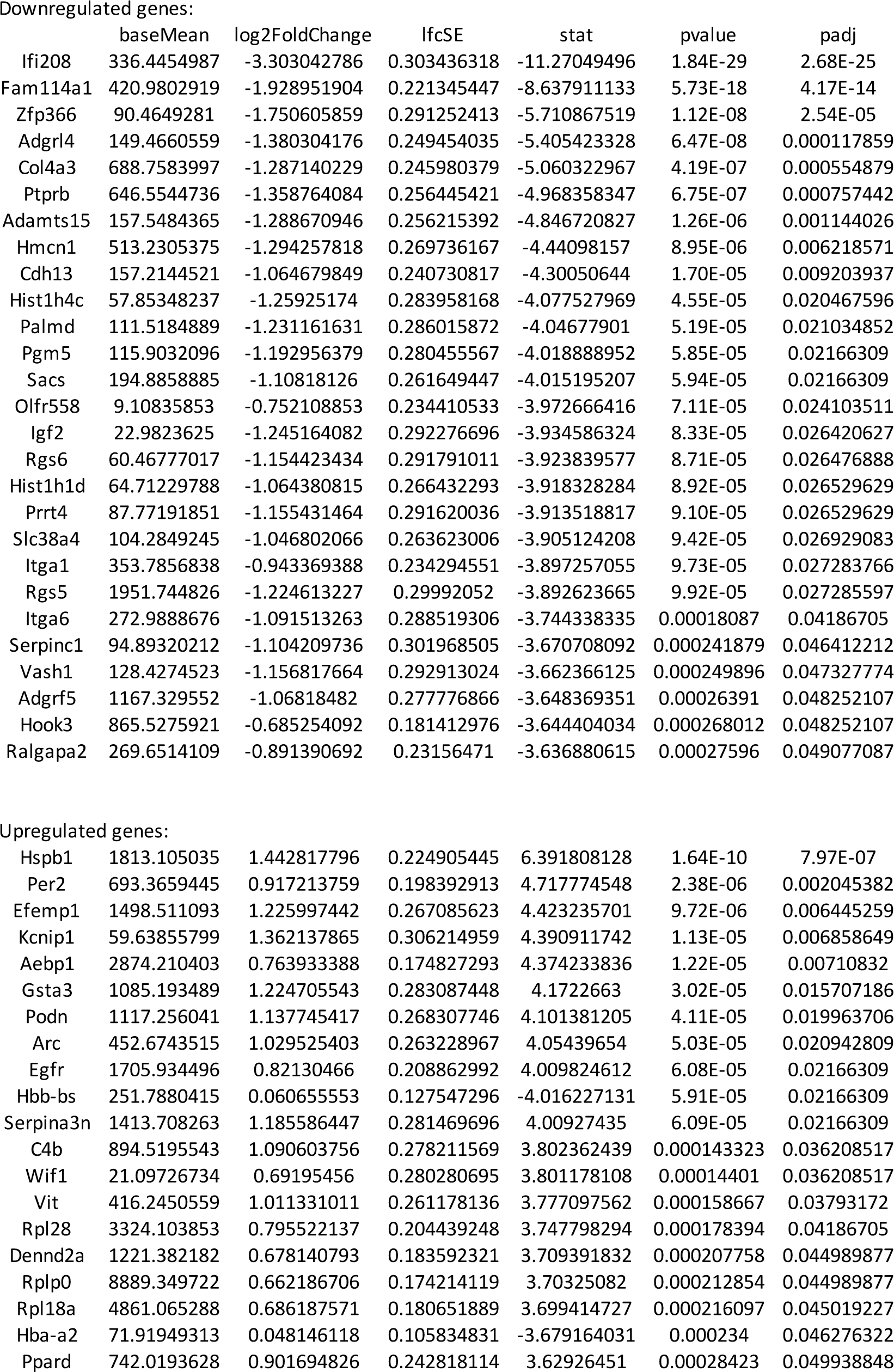
Differentially expressed genes in Fam114a1 null cardiac fibroblasts compared to WT CF cells identified by RNA-Seq.

## Notes

### Competing Interest Statement

The authors have declared no competing interest.

